# Cryo-EM of cardiac AL-224L amyloid reveals shared features in λ6 light chain fibril folds

**DOI:** 10.1101/2025.06.25.661559

**Authors:** Chad W. Hicks, Tatiana Prokaeva, Brian Spencer, Shobini Jayaraman, Noorul Huda, Sherry Wong, Hui Chen, Vaishali Sanchorawala, Francesca Lavatelli, Olga Gursky

## Abstract

In amyloid light chain (AL) amyloidosis, aberrant monoclonal antibody light chains (LCs) deposit in vital organs causing organ damage. Each AL patient features a unique LC. Previous cryogenic electron microscopy (cryo-EM) studies revealed different amyloid structures in different AL patients. How LC mutations influence amyloid structures remains unclear. We report a cryo-EM structure of cardiac AL-224L amyloid (2.92 Å resolution) from λ6-LC family, which is overrepresented in amyloidosis. Comparison with λ6-LC structures from two other patients reveals similarities in amyloid folds. Mutation-induced structural differences in AL-224L include altered C-terminal conformation with an exposed ligand-binding surface; an enlarged hydrophilic pore with orphan density; and altered steric zipper registry with backbone flipping, which likely represent general adaptive mechanisms in amyloids. The results suggest shared features in λ6-LC amyloid folds and reveal how mutation-induced structural changes influence amyloid-ligand interactions in a patient-specific manner.

## INTRODUCTION

AL amyloidosis accounts for nearly ¾ of all systemic amyloid diseases. This debilitating life-threatening systemic disorder is caused by the deposition of monoclonal immunoglobulin LCs as extracellular amyloid fibrils in vital organs, particularly heart and kidney^1^. Annually, the disease afflicts 10-12 new patients per million^2,3^. Cardiac AL amyloidosis is detrimental to patients’ prognosis and is lethal if untreated. Current treatments involve chemotherapy and bone marrow transplantation to eliminate the abnormally proliferating B-cell clone that overproduces the aberrant LC protein, along with proteasome inhibitors, immunomodulatory agents, and monoclonal antibodies to eliminate the LC^2–4^. Transplantation of the afflicted organ (kidney, heart) and supportive treatments are also available to extend the patient’s life and improve the quality of life. However, these treatments are not suited for all patients. New therapies are being developed, including small-molecule LC stabilizers and CRISPR-based approaches targeting the aberrant LC generation^3–5^. These efforts are complicated by the extreme variability of antibodies stemming from gene recombination and somatic hypermutations; as a result, each amyloidosis patient features a unique fibril-forming LC^6–8^. Since the pathological role of fibrils in AL and other forms of systemic amyloidoses is well established^4^, understanding molecular underpinnings for amyloid fibril formation, structure and biological properties may help guide the development of much-needed amyloid-targeting therapies.

In the normal polyclonal LC repertoire, κ-LC and λ-LC proteins are found at a 2:1 ratio, yet AL amyloidosis is most frequently associated with the overexpression of λ-LCs^9^. Moreover, *IGLV6-57* gene segment encoding the variable domain (V_L_) of λ6-LCs is expressed in ∼2% of the polyclonal λ-LC repertoire but accounts for ∼25% of AL amyloid cases^10,11^. This study is focused on the λ6-LC protein subfamily.

Native LC proteins (∼25 kDa) share a common two-domain architecture consisting of the N-terminal variable (V_L_) and C-terminal constant (C_L_) domains joined by a short flexible linker (J-region); each domain acquires a β-sandwich immunoglobulin fold. AL deposits always contain V_L_, either as a full-length LC or LC fragments^12,13^. Most LC mutations in AL patients cluster in V_L_ in or near three antigen-binding loops, called complementarity-determining regions (CDRs). Amyloidogenic mutations have been reported to destabilize the native structure in V_L_, to perturb the protective domain-domain interactions in LC monomers or dimers, to increase protein dynamics/exposure in sensitive regions (including the interdomain linker, the amyloid-promoting regions (APRs) in V_L_, and the proteolysis-prone segments in C_L_), or to stabilize the amyloid structure^5,14–21^. However, oftentimes it is unclear how amyloidogenic mutations promote the misfolding of native LCs to amyloid structure. Moreover, the major triggers of misfolding are different for different LCs^22,23^, suggesting different amyloidogenic pathways for different patients.

This idea is supported by the cryo-EM structural studies; 12 structures of tissue-derived amyloid fibrils formed by LCs from λ1, λ3, and λ6 sub-families are currently available^13,18,21,23–26^. The amyloid folds in these fibrils drastically differ from the native immunoglobulin fold. Importantly, fibrils from the same organ of different patients show different amyloid structures^23^, while fibrils from different organs of the same patient show similar structures^24^. Since each AL patient features a unique set of mutations in the fibrillar LC, these results suggest that LC mutations are critical for determining amyloid fold.

Nevertheless, cryo-EM structures of several λ3-LCs from different patients show some similarities, suggesting that amyloids from the same protein subfamily can be structurally related^13,23^. Despite this recent progress, the paucity of the AL amyloid fibril structures and their diversity prohibit using machine-learning tools such as AlphaFold^26^ to predict new amyloid structures of LCs based on their amino acid sequences.

Of the available LC fibril structures, only two proteins are from the λ6-LC subfamily: amyloids from the heart and kidney of patient AL55 (AL-Base number AL55, PDB ID: 6HUD and 8CPE)^24,26^ and amyloid from the heart of a patient AL-9ELS (PDB ID: 9ELS)^21^. Cardiac and renal AL55 amyloids have similar structures that differ from the AL-9ELS structure. To expand the available structural repertoire of λ6-LC amyloids, we determined a cryo-EM structure of cardiac AL-224L amyloid (PDB ID: 9OKA). This structure shares key similarities but also shows important mutation-induced structural differences with AL55 and AL-9ELS amyloids. Interestingly, AL-224L amyloid shows an altered C-terminal conformation with an exposed ligand-binding surface that may bind collagen; an enlarged hydrophilic pore with orphan density; and altered steric zipper registry with backbone flipping. These results suggest shared structural features for λ6-LC amyloids, show how AL mutations influence amyloid structure, and reveal how this structural variability alters amyloid-ligand interactions that define the biological properties of amyloids.

## RESULTS

### Analysis of AL-224L amyloid deposits and the extracted fibrils

Amyloid fibrils were extracted from the explanted heart of a previously reported 63-year-old female patient AL-224 with AL amyloidosis^28^ (Table S1). Tissue analysis by Congo red staining and light microscopy indicated amyloid (Fig. S1A,B). Immunogold transmission electron microscopy showed reactivity with λ-LC antibody (Fig. S1C,D). The *IGL* gene sequence derived from the patient’s bone marrow showed that AL-224L protein belonged to the λ6-LC family and was encoded by the *IGLV6-57*01, IGLJ2*02,* and *IGLC2*02* germline genes.

Liquid chromatography tandem mass spectrometry (LC-MS/MS) of fibril extracts confirmed the sequence identity of the amyloid-forming and the bone marrow-derived LCs and showed 74% peptide coverage in AL-224L residues 1-192, which encompass V_L_ and a part of C_L_ domain (Figs. 1A, S2). 2D-PAGE of fibril extracts showed two most abundant species at 15 kDa (Fig. S2A-B). LC-MS/MS of these species detected AL-224L residues 1-116 (Fig. S2C) confirming that the fibrils contained the entire V_L_ domain. Similar to previous reports^12,28,29^, 2D western blot detected additional low-abundance species representing the full-length and fragmented LC (Fig. S2A-B).

**Figure 1.**
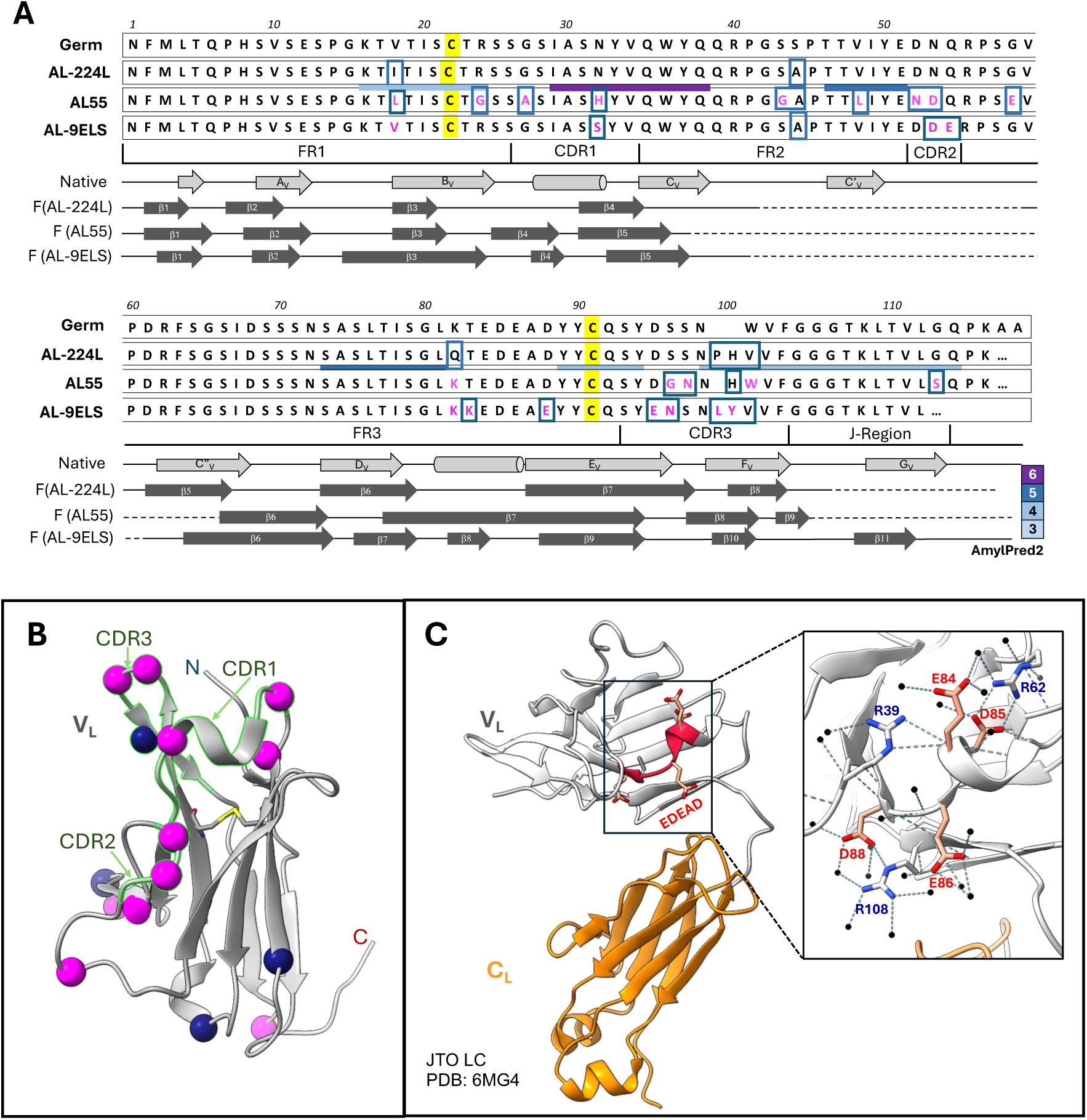
Structural properties of λ6-LCs explored in this study. **A**, Top to bottom: Amino acid sequences of the λ6 germline, AL-224L, AL55, and AL-9ELS VL domains (consecutive residue numbering). AL55 and AL-9ELS residue differences compared to AL-224L (magenta). AL-224L, AL55, and AL-9ELS residue differences compared to germline LC (boxed). Amyloid-promoting regions in AL-224L (horizontal bars) are colored according to the consensus number predicted using AmylPred2^35^. Framework regions (FR), complementarity determining regions (CDRs), the J-region, and the proximal end of CL domain are marked. Linear diagrams show secondary structures of λ6-LCs including β-strands (arrows), α-helices (cylinders), loops/turns (solid lines), and disordered regions (broken lines). “Native” denotes the native fold according to the x-ray crystal structure of a related λ6-LC protein, JTO (PDB ID: 6MG4); “F” denotes amyloid fibril structures of AL-224L (PDB ID: 9OKA), AL55 (PDB ID: 6HUD), and AL-9ELS (PDB ID: 9ELS) determined by cryo-EM. **B**, Native structure of VL domain (PDB ID: 6MGT). CDRs are highlighted. Spheres mark residues differentiating AL55 from AL-224L (magenta) and AL-224L from the germline LC (blue). **C**, Native structure of full-length LC. The acidic-rich segment 84-88 (red). Zoomed-in view of the boxed area shows salt bridges and hydrogen bonds stabilizing the charges in this acidic segment.

LC-MS/MS identified additional proteins in the fibril extracts; Table S2 lists top 30 proteins. AL-224L was prevalent based on normalized spectral abundance factors. Other proteins, including amyloid signature proteins (apoE, apoA-IV) and three chains of collagen VI (ColVI), were identified with lower abundance. ColVI showed similar abundance in prior studies of cardiac λ3-AL59 fibrils that were extracted using similar protocols and contained ColVI bound to LC fibrils^13^.

The native structures of LC and its V_L_ domain (Fig. 1B,C) differ drastically from their structure in an amyloid fibrils^13,18,21,23–26^. To obtain high-resolution structural information on AL-224L fibrils, we used cryo-EM (Fig. S3). While cryo-EM data of amyloid fibrils are typically processed using Relion^30^, we performed cryo-EM data processing exclusively in CryoSPARC. Only two other investigations have determined amyloid structures using CryoSPARC^31,32^. To our knowledge, our structure of AL-224L amyloid has the highest global resolution of all similar structures processed in CryoSPARC. Methods provide detailed information of our data processing workflow to assist future investigations.

Cryo-EM micrographs showed long twisted AL-224L fibrils with either single- or double-filament morphology (Fig. 2A). 2D classification showed that ∼75% of the fibril segments were single-filament with a helical crossover of approximately 700 Å, while ∼25% were double-filament (Fig. 2B). We determined the cryo-EM structure of the single-filament polymorph to a resolution of 2.92 Å (Table 1 and Fig. S4A), which showed clear side-chain densities (Fig. S4B-D) and β-sheet separation (Fig. S4E). Local resolution estimation of the cryo-EM map showed a well-resolved central core at ∼2.5 Å and less resolved peripheral parts at ∼3.5 Å (Fig. 2C). Symmetry search analysis of the single-filament structure revealed a helical rise of 4.86 Å and a twist of 1.20°. Analysis of the corrugation in the EM density of backbone carbonyl atoms suggested a right-handed twist (Fig. S5). We were unable to determine the double-filament structure, likely due to an insufficient amount of data for this polymorph.

**Figure 2.**
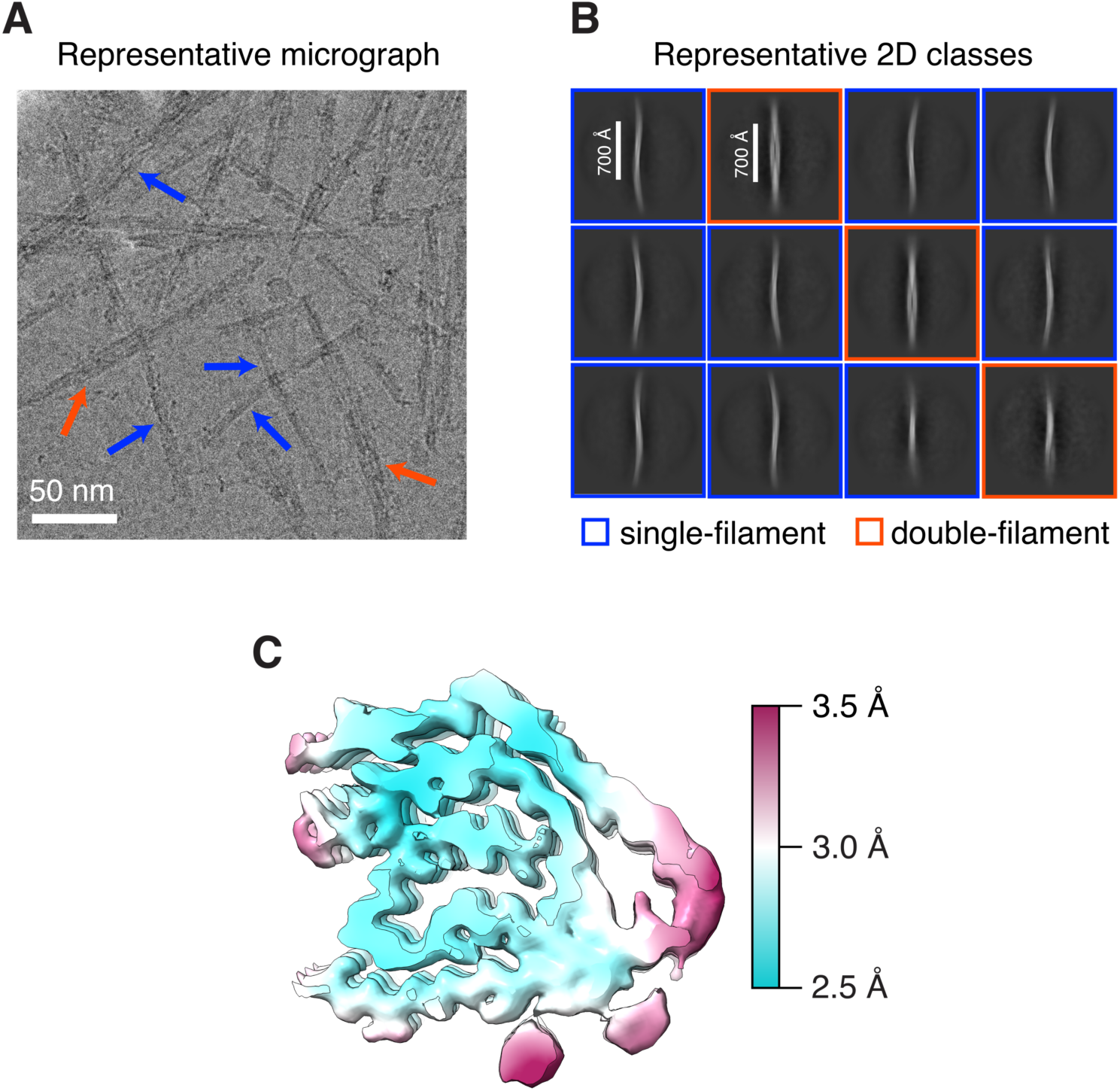
Amyloid fibril polymorphs and local resolution of the single-filament AL-224L structure. **A**, Representative cropped micrograph shows single (blue arrows) and double filaments (red arrows). **B**, Representative 2D classes at a box size 1800 px (1620 Å) show an approximate crossover distance of 700 Å for both single and double filaments. **C**, Cross-section of one molecular layer of the cryo-EM density map colored by local resolution.

**Table 1.** Cryo-EM statistics and validation metrics. Table of details related the cryo-EM structure of AL-224L amyloid. The details include parameters for data collection, data processing, model building, and validation.

| Parameter | Data Collection |
| --- | --- |
| Magnification (X) | 130,000 |
| Voltage (kV) | 200 |
| Electron exposure ($e^-/\text{\AA}^2$) | 40 |
| Dose rate ($e^-/\text{px/s}$) | 10.82 |
| Defocus range ($\mu\text{M}$) | -0.8 to -2.5 |
| Pixel size ($\text{\AA}$) | 0.900 |
| Camera | Falcon 4i |
| Energy Filter slit width (eV) | 10 |
| Micrographs (#) | 10,309 |
|  | **Data Processing** |
| Box size (px) | 400 |
| Segment separation distance ($\text{\AA}$) | 45 |
| Initial uncleaned segment stack (#) | 421,024 |
| Helical rise ($\text{\AA}$) | 4.86 |
| Helical twist ($^\circ$) | 1.20 |
| Particle duplication symmetry | C1 |
| Final segments (#) | 214,100 |
| Map resolution at 0.143 FSC cutoff ($\text{\AA}$) | 2.92 |
|  | **Model Building and Validation** |
| Non-hydrogen atoms (#) | 3,335 |
| Subunits | 5 |
| CC mask | 0.75 |
| Bond length RMSD ($\text{\AA}$) | 0.002 |
| Bond angles RMSD ($^\circ$) | 0.602 |
| Clash score | 7.82 |
| Ramachandran Outliers (%) | 0 |
| Ramachandran Allowed (%) | 6.02 |
| Ramachandran Favored (%) | 93.98 |
| Rotamer Outliers (%) | 0 |
| C-beta outliers (%) | 0 |
| MolProbity score | 1.83 |

### Cardiac λ6-LC amyloids from different patients share key structural features

Like in other patient-derived AL fibrils, the fibril core in AL-224L contains misfolded V_L_ domains in a parallel in-registry cross-β-sheet conformation (Fig. 3). The β-strands are linked via intermolecular hydrogen bonds running along the fibril z-axis while the side chains pack via steric zippers in the x-y plane^33,34^.

**Figure 3.**
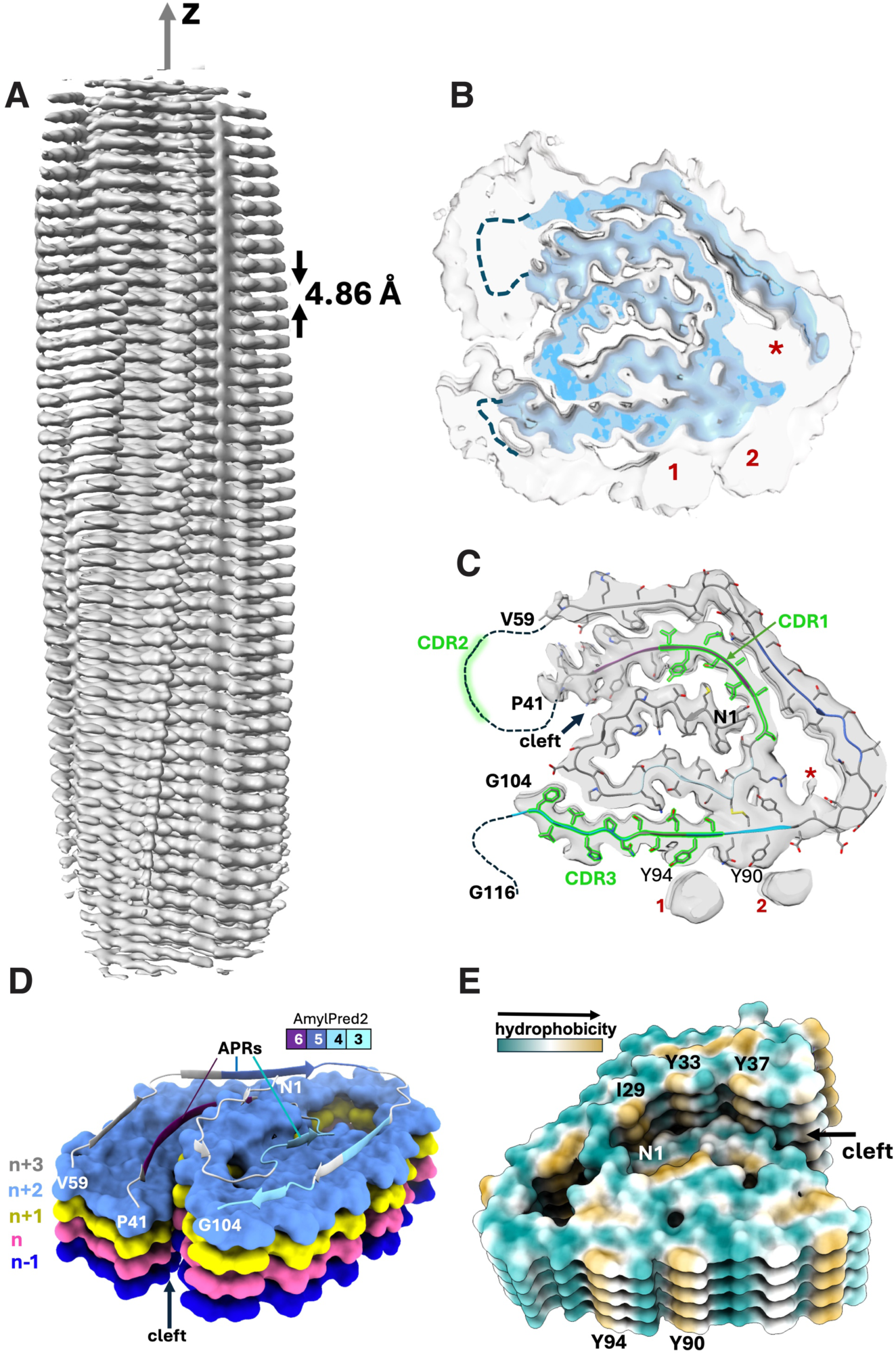
Structural overview of the AL-224L single-filament amyloid polymorph. **A**, Sharpened cryo-EM map (2.9 Å) of AL-224L amyloid (side view). **B**, EM map (top view) showing well-ordered fibril core (blue, contouring threshold 0.0994) and lower-order regions (gray, threshold 0.0325). Disordered segments 42-58 and 105-116 (dashed lines). **C**, Cross-section of one fibril layer superimposed on the cryo-EM map. Stick model shows ordered segments N1-P41 (NT-segment) and V59-G104 (CT-segment). CDRs are colored in green and orphan densities are labeled as 1, 2, and *. **D**, A stack of molecular layers shows n to n+1 and n to n+2 interlayer contacts across the central cleft (top view). APRs in the top layer are color-coded from least amyloidogenic (cyan) to most amyloidogenic (purple), as in Fig. 1A. **E**, Surface representation of five molecular layers colored according to side chain hydrophobicity (bottom view).

The AL-224L V_L_ domain contains 111 amino acids, while its AL55 counterpart contains 110 amino acids that differ from AL-224L in 13 point substitutions and 1 insertion (Fig. 1A). Surprisingly, the two amyloids show similar folds consisting of a central N-terminal “snail shell-shaped” segment^26^ (NT-segment, residues 1-41 in AL-224L and 1-34 in AL55) enclosed by a C-terminal C-shaped segment (CT-segment, residues 59-104 in AL-224L and 66-106 in AL55) (Fig. 4A,B). These structured antiparallel segments are covalently linked by the canonical disulfide, C22-C91. This ordered structure is flanked by two disordered segments seen in the cryo-EM map as diffuse densities, one encompassing the CDR2-containing linker between the NT- and CT-segments (residues 42-58 in AL-224L and 35-65 in AL55) and the other extending C-terminally from V_L_ into the proximal end of C_L_ (residues 105-116 in AL-224L and 107-119 in AL55) (Figs. 3B,C, 4A,B). The ordered C-terminal tail (CT-tail, residues 95-104 in AL-224L and 95-106 in AL55) is repositioned: the AL-224L CT-tail extends straight around the NT-segment while the AL55 CT-tail makes two ∼90° turns and folds back on itself (Fig. 4A,B).

**Figure 4.**
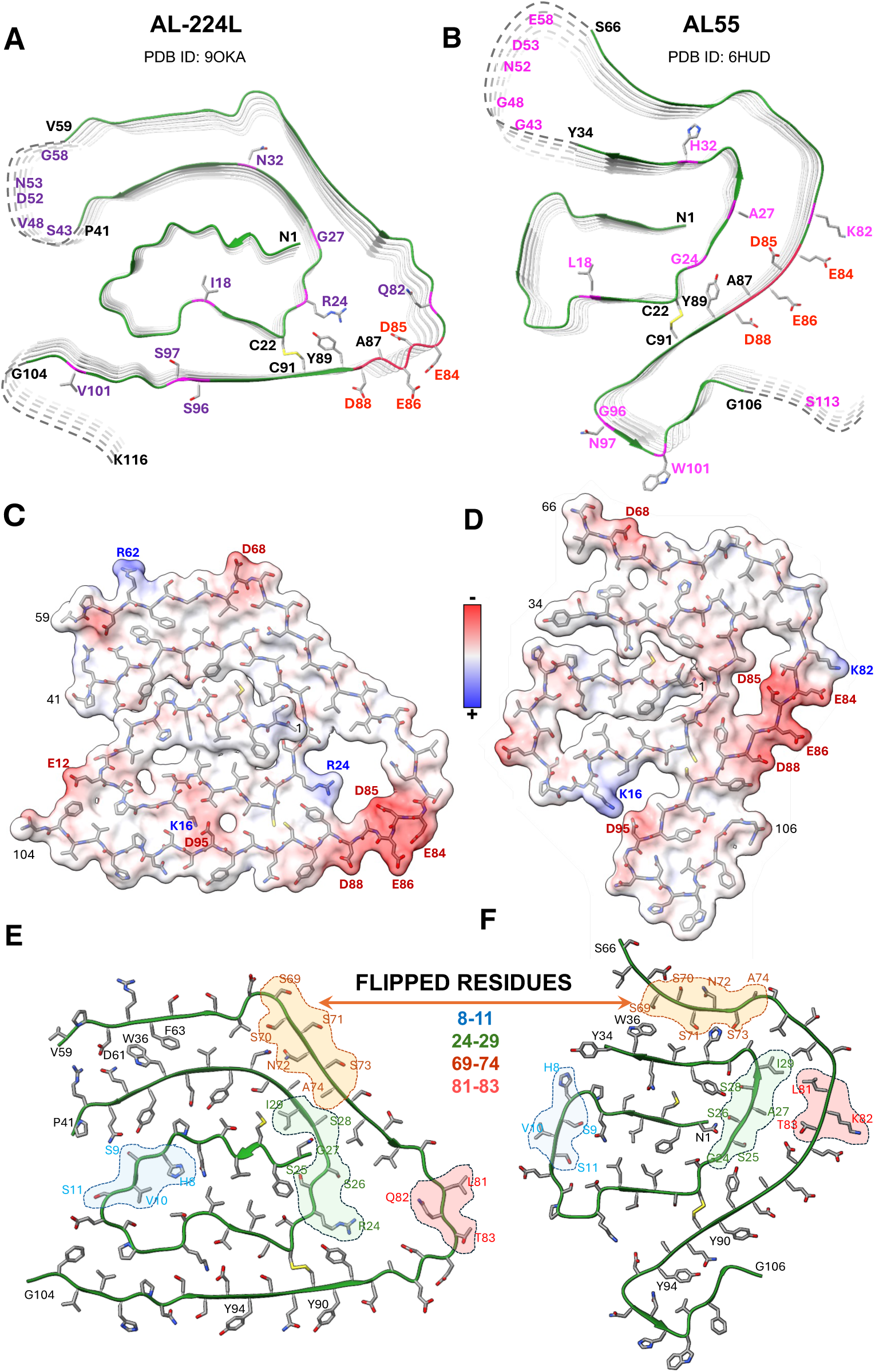
Comparison of AL-224L and AL55 fibril structures. **A, B**, Cross-sectional top views show proteins’ backbone in five molecular layers. Residues that differ in AL-224L (violet) and AL55 (magenta), residues in the acidic-rich segment (red), and other residues mentioned in the manuscript (black) are shown. **C, D**, Space-filling model of one fibril layer colored according to coulombic potential. **E, F**, Regions showing opposite side chain orientation (backbone flipping) in AL-224L vs. AL55 amyloids.

The AL-9ELS V_L_ domain contains 111 amino acids^21^ that differ from AL-224L in 11 point substitutions (Fig. 1A). Like AL-224L and AL-55, the AL-9ELS amyloid structure contains antiparallel well-ordered NT- and CT-segments (residues 1-40 and 61-112) connected by the disordered CDR2-containing linker and by the C22-C91 disulfide (Fig. 5). The NT-segment in the AL-9ELS amyloid is not snail-shell shaped, like in AL-224L and AL55. Rather than occupying a sequestered central position, AL-9ELS segment 1-14 is repositioned towards the structural periphery and rotates by ∼180° to encircle residues 96-101 of the CT-segment (Fig. 5C). In contrast with AL-224L, the AL-9ELS CT-tail (structured residues 95-112) folds back on itself.

**Figure 5.**
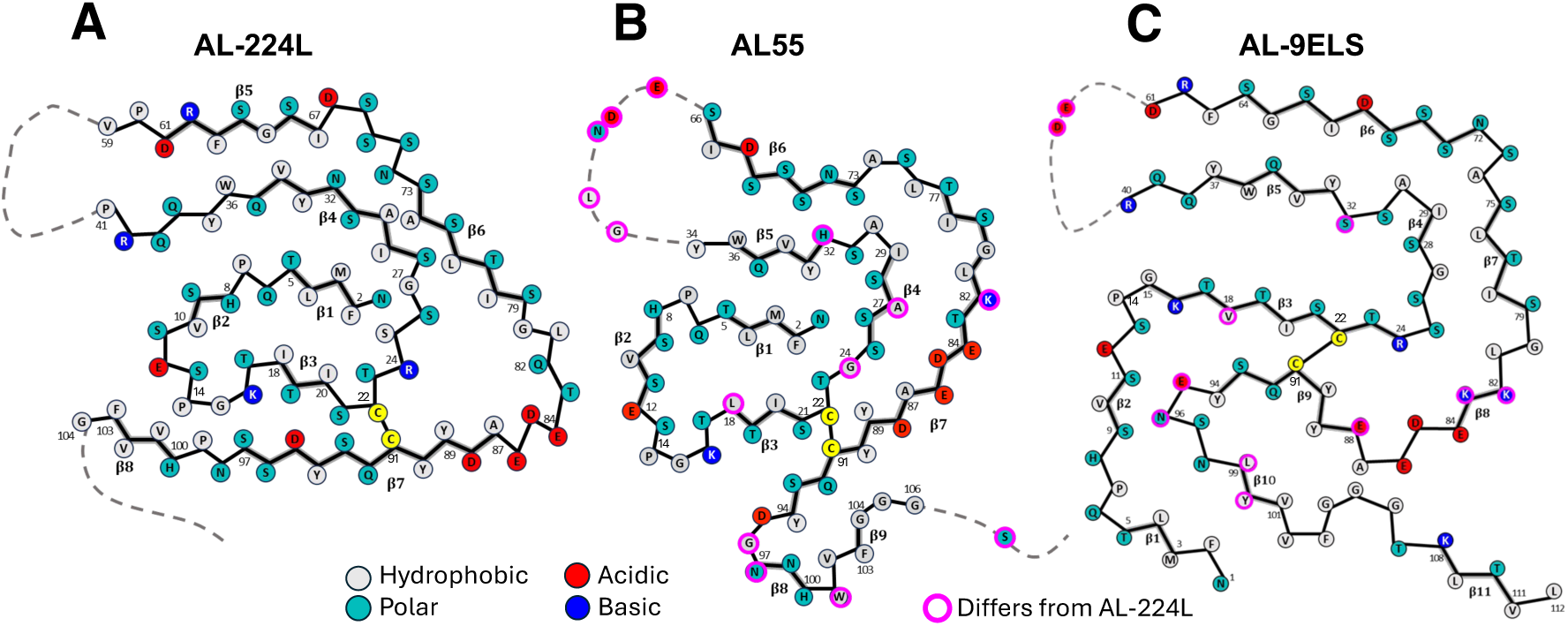
Three available λ6-LC cardiac amyloid structures. Schematic diagram showing the polypeptide sequence and fold in the fibril core of AL-224L (PDB ID: 9OKA) (**A**), AL-55 (PDB ID: 6HUD) (**B**), and AL-9ELS (PDB ID: 9ELS) (**C**). β-strands shown as thick lines; disordered regions shown as dashed lines.

All three λ6-LC amyloids contain a buried salt bridge, K16-D95 (E95 in AL-9ELS) (Figs. 4C,D, 5)^21,26^. This stabilizing salt bridge and the C22-C91 disulfide help maintain the antiparallel orientation and local registry of the juxtaposed NT- and CT-segments.

V_L_ domain of LCs typically harbors six amyloid-promoting regions (APRs)^19^. Six APRs were predicted by at least three different methods for AL-224L (Fig. 1A) and other λ6-LCs using the sequence-based consensus algorithm AmylPred2^35^. Four APRs are in the structured segments of AL224L, AL55, and AL-9ELS fibrils (Figs. 1A, 3D); one APR overlapping CDR2 is disordered; and one APR overlapping CDR3 acquires variable conformations in different amyloids^19,21^. The disulfide-forming residues C22 and C91 are located in APRs in the structured regions near CDR1 and CDR3 (Fig. 1A), supporting the importance of the disulfide-containing regions in amyloid formation^21,36^.

The structured segments in AL-224L and AL55 amyloids are flattened but not planar and are stacked in β-helix-like structures. In these structures, the rugged fibril ends expose hydrophobic side chains I29, Y33, and Y37 from the major APR (Fig. 3E), which may help recruit new protein molecules to the growing fibril end^26^. Residue segment N1-P14 protrudes from the plane of the CT-segment and interacts with strands across the central cleft that are one or two molecular layers up the fibril length, n to n+1 and n to n+2 (Fig. 3D). AL-9ELS amyloid also shows cross-layer interactions involving N1-P14 segment^21^. In all three amyloids, such cross-layer interactions sterically interlock different molecules contributing to fibril stability.

### Mutations in CDR3 alter the CT-segment conformation affecting ligand binding

Mutations in the ordered segments of λ6-LC amyloids, which include CDR1 and CDR3, probably contribute to their structural differences (Fig. 5), while mutations in disordered segments such as the CDR2-containing linker probably influence the amyloid formation kinetics but not necessarily the ordered structure^22^. Amino acid sequence in CDR3 of λ6-LCs is particularly variable^6,7^. Compared to AL-224L, AL55 residues 96-101 from CDR3 differ in three substitutions and one insertion, while CDR3 residues in AL-9ELS differ in four substitutions, resulting in different CT-tail conformations. Importantly, in AL-224L amyloid, the CT-tail extends along the NT-segment exposing the side chains of Y90, Q92, and Y94 (Fig. 4A). However, in AL55 and AL-9ELS amyloids, the CT-tail folds back on the structured CT-segment and occludes these side chains (Figs. 4B, 5B,C).

The Y90 and Y94 rings form one of the most hydrophobic solvent-exposed surfaces on the AL-224L fibril (Fig. 3E), which is energetically unfavorable. Our cryo-EM map shows two large orphan densities, one near Y94 and Q92 side chains and another near Y90 and D88 side chains (Figs. 3C, 6A,B). To test the significance of these densities, we filtered the unmasked map to a high statistical significance threshold cutoff using false-discovery rate thresholding^37^ (Supplemental Figure 1). The orphan densities were visible at a high threshold cutoff, excluding the possibility that they were noise and suggesting bound ligands. These ligands are absent from AL55 and AL-9ELS structures where this fibril surface is occluded by the CT-tail^21,26^ (Fig. 5B,C). These results show how high sequence variability in CDR3 can translate into structural variability in the λ6-LC amyloid, leading to distinct patient-specific CT-tail conformations that critically influence amyloid-ligand binding.

**Figure 6.**
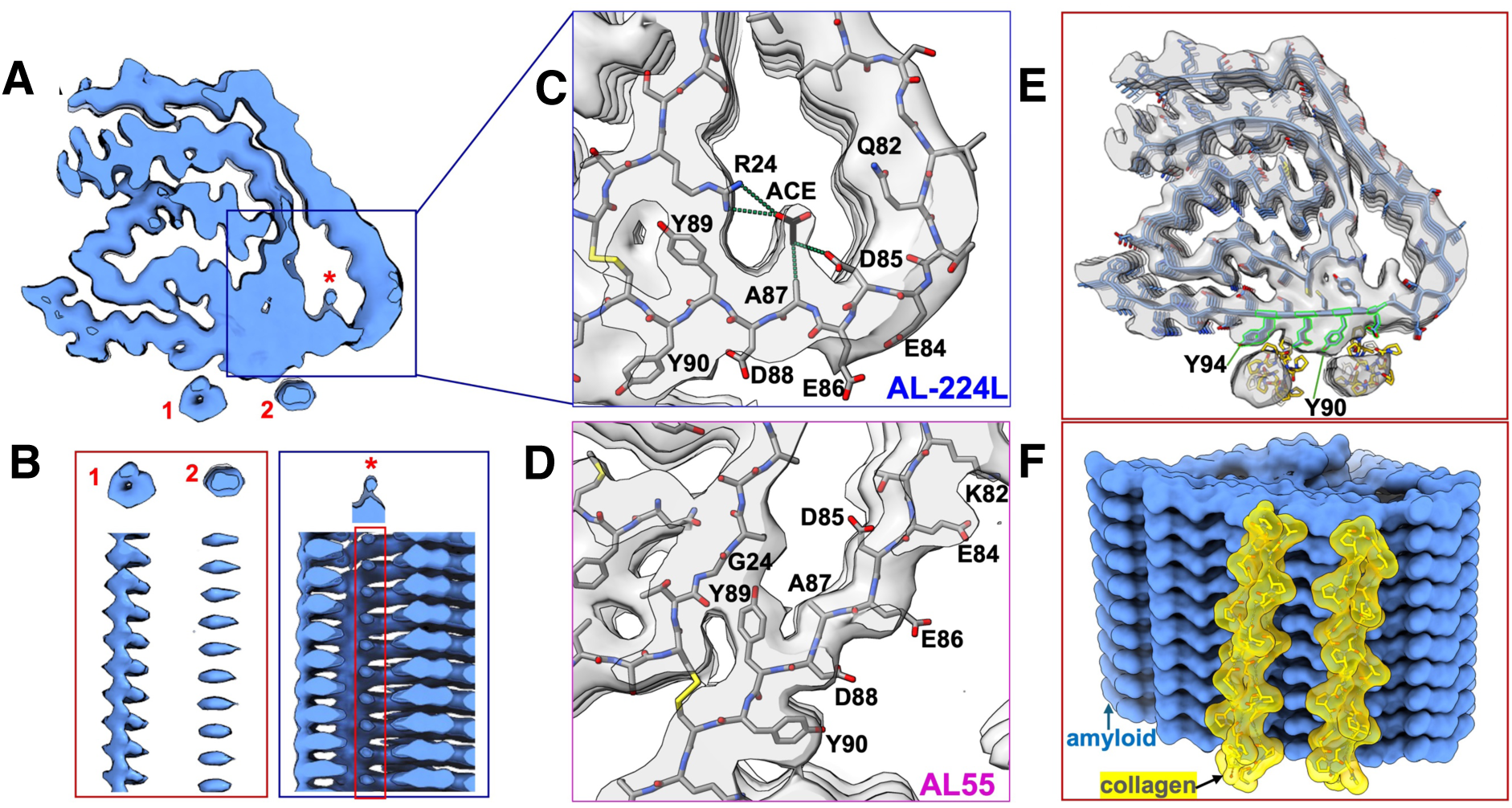
Orphan densities in the cryo-EM map of AL-224L amyloid. **A**, Sharpened cryo-EM density map shows one fibril layer (top view) with orphan densities at external (1, 2) and internal (*) sites. **B**, Side views show orphan densities in the maps at a threshold of 0.0781 (1, 2) or 0.0567 (*). **C**, Zoomed-in view of the EM map and model of the hydrophilic pore in AL-224L shows acetate (ACE) molecule manually fitted in the extra density. Dotted lines show acetate-protein contacts within 4 Å. **D**, A view similar to panel C of the amyloid pore in AL55. **E**, Cryo-EM map and model of AL-224L show two top docking poses of collagen triple helix (PDB ID: 1CAG). **F**, Side view of the space-filling model shows top two docking poses of collagen (yellow) to AL-224L amyloid (blue).

### AL-224L shows orphan density in a hydrophilic pore near an acidic-rich segment

Uncompensated charged segments in amyloids are highly energetically unfavorable due to a strong repulsion within and between molecular layers^38,39^. Conflicting interatomic forces in such segments can generate structurally frustrated regions with variable conformations^39^. LCs feature a highly charged conserved residue segment 84-88 (EDEAD in AL-224L and AL55, EDEAE in AL-9ELS). In the native LC structure, this segment is stabilized by an extensive network of salt bridges and hydrogen bonds (Fig. 1C). In λ6-LC amyloids, the acidic segment is located at the fibril surface forming a part of a variable hydrophilic pore (Fig. 5). In AL-224L amyloid this pore is enlarged and the EDEAD segment shows reduced local resolution indicating structural frustration (Fig. 2C).

Mutations in and near the pore probably contribute to differences in its structure. These differences between AL-224L and AL55 include R24G, G27A, and Q82K substitutions. In the narrow pore of AL55, the Y89 ring packs against G24 (Fig. 4B), but in AL-224L amyloid, the Y89 side chain orients towards the C22-C91 disulfide and forms a cation-π interaction with R24 (Fig. 4A). In AL-9ELS amyloid, R24 also interacts with Y89^21^. Notably, Y89 is highly conserved in λ6-LCs^11^. Therefore, R24G substitution alters the conformation of the highly conserved Y89 residue in the pore.

In AL55 and AL-9ELS amyloids the charges within the acidic-rich segment are partially balanced by the K82-E84 salt bridge; additional charge-charge interactions stabilize the acidic-rich segment in AL-9ELS (Fig. 5C). However, in AL-224L amyloid the K82Q substitution eliminates the sole stabilizing salt bridge thus increasing the charge imbalance at the acidic-reach segment (Fig. 4C), destabilizing the local backbone conformation (Fig. 2C), and contributing to the pore enlargement.

The cryo-EM map of AL-224L shows a strong orphan density in the pore near the side chains of R24, A87, and D85 (Figs. 3C, 6A-C). This density is well above the noise level based on the false-discovery rate map thresholding (Supplemental Fig. 1). The cryo-EM study of AL-9ELS also reports an orphan density in the pore near R24 and Y89 side chains^21^. In contrast, AL55 (which contains G24 instead of the bulky R24) shows no orphan density in the pore (Fig. 6D), suggesting that this density is associated with R24 but not G24 λ6-LC variants.

The orphan density in the hydrophilic pore of AL-224L amyloid likely represents a small water-soluble molecule that interacts via its acidic and hydrophobic moieties with R24 and A87, respectively. One possibility is acetate, an abundant tissue metabolite whose cardiac uptake increases in heart failure^40^, which was the cause of death of patient AL-224. Analysis of fibril tissue extracts using an ELISA-based assay detected short-chain fatty acids (Table S4 and Fig. S6) confirming them as potential candidates for binding amyloid in tissues. Figure 6C illustrates how an acetate molecule can fit in the orphan density in the pore of AL-224L amyloid.

### Registry shifts and backbone flipping in amyloid helps accommodate AL mutations

Compared to AL55, the wider pore in AL-224L amyloid is facilitated by a shift in the backbone registry of the steric zippers between the CT- and NT-segments (Fig. 4E,F). This registry shift propagates from the EDEAD segment towards the CDR2-containing linker encompassing a 30-residue segment 59-88. For example, in AL-224L the W36 side chain packs between the D61 and F63 side chains, yet in AL55 the W36 side chain packs against S69 (Fig. 4E,F). To accommodate such a 7-residue registry shift, the backbone in segment 69-74 is flipped in AL-224L relative to AL55, inverting the side chain orientation in this segment. Registry shifts with the backbone flipping are also seen in other structured residue segments including 8-11, 24-29, 69-74, and 81-83 (Fig. 4E,F). As a result, most residues pack differently in the X-Y plane of AL-224L and AL55 amyloids, yet the overall amyloid fold is largely preserved.

The AL-9ELS structure also shows registry shifts with backbone flipping. For example, segment 32-40, which packs antiparallel to segment 61-70 with one-residue registry shift between AL-9ELS and AL-224L amyloids, shows an opposite side chain orientation in AL-9ELS compared to both AL-224L and AL55 amyloids (Fig. 5). These findings suggest that registry shifts and associated backbone flipping is a plausible general mechanism enabling amyloids adapt to mutations while largely preserving the amyloid fold (Fig. 7).

**Figure 7.**
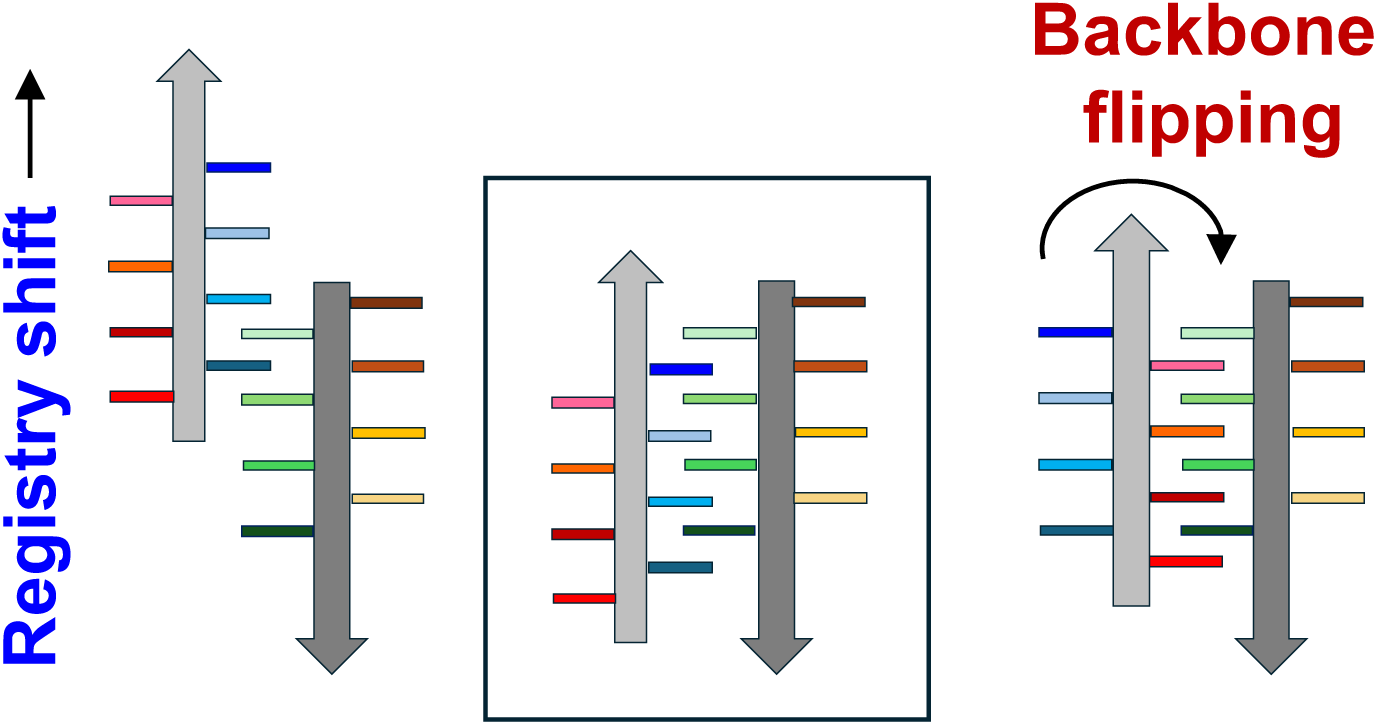
Schematic diagram illustrating registry shifts and backbone flipping in a steric zipper. A steric zipper before registry shift and backbone flipping (middle). A steric zipper after registry shift (left). A steric zipper after backbone flipping (right). b-strands are shown as gray arrows; side chains are shown as colored bars.

## DISCUSSION

### λ6-LC amyloids have shared structural features

We determined the single-filament structure of AL-224L cardiac amyloid at 2.92 Å resolution and compared it with the available amyloid structures of two other cardiac λ6-LCs, AL55 (4.0 Å resolution) and AL-9ELS (3.0 Å resolution). Despite 14 amino acid differences between AL-224L and AL55, these two amyloids have surprisingly similar overall folds (Fig. 4), while all three amyloids share key structural features (Fig. 5). These features include a structured fibril core comprised of antiparallel NT- and CT-segments connected by a disordered CDR2-containing linker. In λ1-LC and λ3-LC amyloids the CDR2-containing linker can be either ordered or disordered^13,23^. Importantly, in all LC amyloids NT- and CT-segments are linked by the conserved internal disulfide that maintains their antiparallel orientation providing a major constraint for amyloid formation and structure^13,21,23,25^. Furthermore, in λ6-LC amyloids the planar part of the CT-segment partially encircles the fibril surface, while the non-planar NT-segment forms cross-layer interactions. Lastly, the amyloid structure adjacent to the C22-C91 disulfide contains a variable pore; the pore is lined by mostly hydrophilic residues including the acidic-rich segment 84-88 (Fig. 4C,D), while the Y89 side chain packs against G24/R24 in the pore (Figs. 4A,B). Since in λ6-LCs residues 85-91 are highly conserved while residue 24 is typically either G or R^7,11^, these structural features probably extend to other λ6-LC amyloids.

The observed structural similarities between the λ6-LC amyloids are facilitated, in part, by the registry shifts with backbone flipping in steric zippers (Figs. 4E,F, 7). Importantly, we were able to attribute the structural differences between amyloids to LC mutations, providing clear insights into the role of patient-specific AL mutations in amyloid polymorphism and ligand binding.

### Germline λ6-LC is compatible with the amyloid folds in tissue-derived fibrils

The fibril-forming V_L_ domain in λ6-LCs is encoded by the germline gene *IGLV6-57* ^10,11^. We propose that the hypothetical protein with the germline amino acid sequence can be accommodated in the structure of AL-224L and/or other tissue-derived amyloids (Fig. 1A). The germline-encoded residue V18 (I in AL-224L, L in AL55, and V in AL-9ELS) can be readily accommodated instead of I18 of AL-224L in the structured NT-segment in the hydrophobic amyloid core. The germline-encoded S44 (A in AL-224L, AL55, and AL-9ELS) can also be accommodated in the flexible CDR2-containing segment. Furthermore, the germline-encoded residue K82 (Q in AL-224L, K in AL55 and AL-9ELS) forms a stabilizing salt bridge with E84 in AL55 and AL-9ELS amyloids (Figs. 4D, 5B,C)^21,26^. Lastly, germline mutations in positions 99-101 of CDR3 are very different in AL-224L, AL55, and AL-9ELS (Fig. 1A) and probably influence the variable CT-tail conformation at the periphery of the amyloid structure (Fig. 5). This analysis suggests that the amyloid folds observed in AL-224L, AL55, and AL-9ELS are compatible with the germline LC, suggesting that aspects of these folds are probably shared by other λ6-LC amyloids.

The amyloid structure of AL-9ELS differs significantly from those of AL-224L and AL55. The major difference involves residue segment 1-14 that forms cross-layer interactions in all three amyloids. In AL-224L and AL55 amyloids, residues 1-14 are centered in the snail shell-shaped core (Fig. 5A,B), yet in AL-9ELS these residues wrap around the structural periphery (Fig. 5C)^21^. Since residues 1-14 are identical in the three λ6-LCs (Fig. 1), their conformational differences must reflect mutations located elsewhere. The likely candidates include residues 18 and 32 in the middle of the snail shell-shaped NT-segment (which harbors segment 1-7 in AL-224L and AL55 amyloids) and residues 95, 96, 99 and 100 in the CT-tail (which interacts with residues 1-14 in AL-9ELS amyloid) (Fig. 5). This comparison illustrates how the combined effects of several AL mutations define the amyloid conformation in a mutation-free segment.

### Low predicted stability of λ6-LC amyloid fibrils may promote pathology

Prior biophysical studies of the natively folded recombinant human λ6-LCs did not show reduced thermodynamic stability of AL-224L compared to non-amyloidogenic controls but revealed subtle differences in local stability and proteolytic processing, which were relevant to fibrillogenesis^19^. In the current study, we assessed the amyloid fibril stability. PDBePISA software^41^ was used to calculate free energy change upon exposure of one fibril layer to solvent for AL-224L and AL55 amyloids, as well as for the hypothetical germline λ6-LC sequence placed in these fibril structures.

AL-224L has slightly weaker solvation energy than AL55, indicating that it is slightly less stable than AL55 (Table S3). Structures of AL-224L or AL55 amyloids containing the hypothetical protein with the germline λ6-LC sequence showed comparable solvation energies as tissue-derived AL-224L and AL55, indicating comparable stabilities (Table S3). These calculations support our hypothesis that the germline λ6-LC sequence is compatible with amyloid structures of the tissue-derived fibrils.

Next, we computed solvation energies for all other available LC amyloid structures using PDBePISA. AL-224L and AL55 amyloids has lower solvation energies than λ1-LC and λ3-LC amyloids (Table S3). The Saelices team also reported lower solvation energies for AL55 and AL-9ELS compared to λ1-LC and λ3-LC amyloids^21^. These results suggest that, compared to other amyloids from the λ-LC family, λ6-LC fibrils are relatively unstable and can be fragmented more easily, potentially facilitating amyloid proliferation by seeding and enhancing amyloid cytotoxicity^42^.

### Role of registry shifts with backbone flipping in amyloid structures

Comparison of AL-224L, AL55, and AL-9ELS amyloid structures revealed a shift in the steric zipper registry with the backbone flipping in several well-ordered segments (Figs. 4E,F, 5). We posit that such amyloid-specific structural changes provide a general mechanism for adapting the fibril core to protein mutations, post-translational modifications, and structurally frustrated regions that are often interspersed with β-strands in amyloids^39^. Registry shifts and backbone flipping in steric zippers help explain why the risk of amyloid formation increases for proteins containing low-complexity domains and amino acid repeats, such as poly-Q motif in huntingtin or tandem repeats in bacterial amyloids^43–45^. Such repeating units can facilitate formation of steric zippers differing in the backbone registry and side chain orientation to generate energetically equivalent but conformationally distinct fibril polymorphs of the same or closely related proteins^46^.

### A conserved acidic-rich segment is a potential gatekeeper in LC amyloid formation

Unbalanced charges in amyloids are highly energetically unfavorable due to a strong repulsion among molecular layers of parallel in-registry cross-β-sheets^38,39^. LCs contain a conserved residue segment 85-91^11^ that overlaps the acidic-rich segment 84-88, EDEAD. We hypothesize that this acidic segment, which forms stabilizing interactions in the native structure (Fig. 1C) but destabilizes the local structure in amyloid (Fig. 2C), acts as a gatekeeper protecting LCs from amyloid formation. However, AL mutations can balance the charge on this acidic segment to promote fibrillogenesis. Examples include K83 that forms a K83-D85 salt bridge in AL-9ELS amyloid (Fig. 5C). Ligand binding at nearby sites, such as the internal site near A87 and the external site near D88 and Y90 in AL-224L (Fig. 6), can also stabilize the acidic-rich segment 84-88 in amyloid and thereby promote fibrillogenesis.

### R24 but not G24 variant is associated with ligand binding in amyloid pore

Residue 24 is a site of an AL-associated polymorphism in λ6-LCs. R24 is the major variant found in the germline, AL-224L, and AL-9EL LCs^47^. Compared to R24, G24, the second-major variant found in AL55, is overrepresented in amyloidosis and is considered more amyloidogenic^11^. Stabilizing electrostatic interactions involving R24 in the native LC structure were proposed to counteract amyloid formation^47^, but the effects of R24G substitution on the amyloid structure and stability were unknown. Cryo-EM structures of AL-224L, AL-9ELS, and AL55 amyloids show that G24 replacement with R24 induces local structural changes in the hydrophilic pore including the pore enlargement (Figs. 5) likely contributing to reduced amyloidogenicity of the R24 variant. Moreover, the orphan density seen in amyloids near R24 in AL-224L and AL-9ELS, but not near G24 in AL55, represents a bound ligand such as acetate (Fig. 6C). Ligand binding in an internal amyloid pore probably occurs during fibril formation and may critically influence fibrillogenesis, suggesting the direct involvement of residue 24 in this process.

### External ligand binding sites in AL-224L fibrils may harbor collagen triple helices

The cryo-EM map of AL-224L shows two orphan densities, one near residues Y94 and Q92 and another near Y90 and D88 (Fig. 6A,E). Mixed hydrophilic/hydrophobic coordinating residues and the ∼18 Å center-to-center separation between these densities suggest that they potentially represent collagen triple helices (diameter 15 Å) and/or apolipoprotein α-helices (diameter 11-13 Å) bound laterally along the fibril spine, as previously reported for ColVI^13^ and apoE^48^. In the current study, LC-MS/MS of the fibril tissue extracts detected three chains of CoIVI as well as apoE and apoA-IV co-purified with fibrils (Table S2). The orphan density coordinated by Q92 and Y94 appears hollow in the x-y plane (Fig. 6A) consistent with a triple helix. To test this hypothesis, we used pyDock software^49^ for rigid body docking of a canonical collagen triple helix to the AL-224L fibril structure. As a model we used the crystal structure of a 30-residue collagen-like peptide containing consensus sequence repeats Pro-Hyp-Gly (PDB ID: 1CAG)^50^. The top two docking poses show triple helices bound along the Y90 and Y94 side chain amyloid arrays (Fig. 6E,F). These docking poses closely superimpose the two orphan densities in the EM map of AL-224L (Fig. 6E). Therefore, these densities likely represent collagen triple helices bound along the fibril surface. The aromatic rings of amyloid stack against the pyrrolidine rings of collagen. We hypothesize that similar interactions mediate longitudinal binding of ColVI to λ3-AL59 cardiac amyloid^13^; in fact, orphan densities seen in the cryo-EM map of AL59 amyloid near the exposed arrays of Y48 and Y35^13^ overlap the top docking poses of the collagen triple helix and hence may represent bound ColVI.

Direct interactions between collagen and aromatic residues have been reported for globular proteins^51^ including proteins featuring an immunoglobulin fold^52,53^ such as an amyloidogenic LC^54^. Our docking results suggest that ring-stacking interactions also mediate collagen binding to amyloids along the fibril spine. Such a longitudinal binding is expected to influence mechanical and biological properties of amyloid fibrils including amyloid nucleation, growth, fragmentation, interactions with cells including phagocytosis, and the uptake of diagnostic dyes^13,55,56^. Future studies will identify additional details of such amyloid-ligand interactions and their roles in vivo.

## Abbreviations

AL: amyloid light chain
APR: amyloid-promoting region
CDR: complementarity determining region
C_L_: light chain constant domain
cryo-EM: cryogenic electron microscopy
LC: light chain
LC-MS/MS: liquid chromatography tandem mass spectrometry;
V_L_: light chain variable domain.

## Acknowledgements

The authors are grateful to Dr. Esther Bullitt, the director of the Boston University Cryo-EM Core Facility where the data was collected, to Dr. Luca Broggini at the University of Pavia for invaluable advice on the cryo-EM structural analysis of amyloids, and to Mei Chen and Steven Kolakowski at the Harvard Center for Mass Spectrometry for their assistance with the proteomics analyses. We gratefully acknowledge the patient’s family for providing tissue samples for amyloid research.

## Author contributions

C.W.H. prepared the cryo-EM grids, collected cryo-EM data, performed data processing, map interpretation, model building and refinement, and prepared graphics.

T.P. initiated the study, oversaw fibril extraction, LC-MS/MS analysis and immuno-EM, interpreted the results, prepared graphics and contributed to draft writing.

B.S. performed fibril extraction, SDS-PAGE and western blotting, prepared samples for LC-MS/MS, contributed to immuno-EM, and prepared graphics.

S.J. performed biochemical assays, calculated solvation energies, performed molecular docking, and contributed to data analysis and interpretation.

N.H. prepared graphics and contributed to structural interpretation.

S.W. optimized sample preparations.

H.C. performed immuno-EM of the tissues and fibrils.

V.S. contributed to the overall study administration and funding.

F.L. oversaw fibril extraction, performed LC-MS/MS analysis and interpretation, and contributed graphics, draft writing, and funding.

O.G. administered the project, interpreted the results, wrote the first draft, and contributed to funding. C.W.H. and O.G. wrote the paper with input from all authors.

## Data availability

The AL-224L amino acid sequence has been deposited to the AL-base, and the gene sequence, *AL3IGLV6-57*01-IGLJ2*02-IGLC2*02* has been deposited in the GenBank (accession number KY432418). The cryo-EM model and map data have been deposited to the Protein Data Bank (PDB ID: 9OKA, EMDB code 70557). Raw cryo-EM movies have been deposited to the Electron Microscopy Public Image Archive (accession code EMPIAR-12815). The LC-MS/MS proteomics data have been deposited to the ProteomeXchange Consortium via the PRIDE partner repository (dataset identifiers PXD064296).

## METHODS

### Clinical data, sample collection and clinical characterization of patient AL-224

The study was conducted in accordance with the Declaration of Helsinki. Informed consent for the data and sample collection was obtained from the patient at presentation with the approval of the Institutional Review Board at Boston Medical Center. Bone marrow aspirate, clinical information, laboratory data, and post-mortem heart tissue were obtained from the sample biorepository and patient database maintained by the Boston University Amyloidosis Center.

Detailed clinical and laboratory characteristics of patient AL-224 are provided in Table S1 and in^19^. Briefly, a 63-year-old female with treatment naïve AL amyloidosis presented with cardiac and soft tissue involvements; λ LC restriction; circulating clonal IgG and λ free LC (FLC) on serum immunofixation electrophoresis; abnormal FLC κ/λ ratio; and λ FLC on urine immunofixation electrophoresis. The abdominal fat aspirate and endomyocardial and bone marrow biopsies were positive for amyloid by Congo red staining imaged using light microscopy in bright and polarized light.

The amyloid fibril typing by either immunoelectron microscopy or mass spectrometry was not available at the time of patient’s evaluation. The patient passed away of cardiac arrest one month after the initial evaluation. Post-mortem evaluation demonstrated interstitial and vascular amyloid deposition in multiple organs, including the heart. The unfixed post-mortem cardiac tissue was frozen at -80 °C; formalin-fixed paraffin-embedded cardiac tissue block was made and used for further studies.

### Histological and immunoelectron microscopy analyses of cardiac tissue

Post-mortem cardiac tissue was evaluated for amyloid by Congo red staining / light microscopy. Amyloid typing in cardiac tissue was performed using immunogold electron microscopy with antibodies directed against λ and κ LCs as previously described^57^. Briefly, for primary antibodies we used polyclonal rabbit anti-human antibodies against λ and κ LCs (Agilent Tech, Cat. #A019102-2 and #A019302-2). Secondary goat anti-rabbit IgG antibody conjugated to 15 nm gold particles (Ted Pella, Cat. #15727) and goat anti-mouse IgG antibody conjugated to 20 nm gold particles (Ted Pella, Cat. #15753) were used. The micrographs were collected using a JEOL JEM1020 transmission electron microscope.

### Gene sequencing

The *IGL* gene was cloned and sequenced from unselected bone marrow plasma cells as previously described^58^. Once the *IGLV* gene was identified, nucleotide sequence errors introduced by FR1 primers were corrected by additional PCR amplification with 5′ primer for the *IGLV6-57* leader region and a 3′ universal *IGLC* primer. The light chain sequence, AL-224L, was deposited in GenBank (accession number KY432418).

### Amyloid fibril extraction from autopsied cardiac tissue

Fibrils were extracted from the tissue as previously described^26^ using overnight tissue digestion with C. histolyticum collagenase (5 mg/ml in a buffer containing 20 mM Tris, 140 mM NaCl, 2 mM CaCl_2_, pH 8.0), followed by 15 cycles of homogenization in Tris EDTA buffer (20 mM Tris, 140 mM NaCl, 10 mM EDTA, pH 8.0) to remove soluble proteins, and by repeated homogenization of the remaining pellet in 1 ml ultrapure water. The fibril-containing supernatants from the water homogenization cycles were retained. To evaluate the protein yield, each 1 ml fraction was quantified using a Pierce BCA Protein Assay Kit (Thermo Fisher Scientific). The cryo-EM and proteomic analyses were performed using the supernatant from the third water-extraction cycle (water fraction 3), which was visibly cloudy. 2D-PAGE and western blot analyses of amyloid fibril extract 2D SDS-PAGE was performed under denaturing and reducing conditions as previously described^26^. Briefly, proteins from water fraction 3 (200 μg) solubilized in the isoelectric focusing buffer (IEF) were first separated on 11 cm strips with immobilized non-linear pH gradient 3–10 (Bio-Rad) using a Bio-Rad Protean IEF cell, and then on 8–16% polyacrylamide gradient gels (Criterion TGX gels, Bio-Rad). Gels were stained with GelCode Blue Stain Reagent (Thermo Fisher Scientific) and imaged using an ImageQuant LAS 4000 (GE). Spots of interest were excised and the proteins were identified by mass spectrometry as described below. For western blotting, proteins were transferred overnight onto a PVDF membrane (Millipore) using Criterion Blotter (Bio-Rad) and probed with a polyclonal rabbit anti-human λ LCs antibody (Fortis Life Sciences) used at a concentration of 0.125 μg/ml.

### Proteomic analysis of fibril extracts by LC-MS/MS

For proteomic analysis of the extracted fibrils from water fraction 3, 25 μg of proteins were subjected to in-solution digestion with MS-grade trypsin (Thermo Fisher Scientific; enzyme:protein ratio 1:20) overnight at 37 °C. For excised protein spots #15 and #16 from the 2D SDS-PAGE, in-gel digestion was performed as described^26^. Samples were cleaned up with Thermo SPE C18 tips and the LC-MS/MS analysis was conducted using Q Exactive HF-X High Resolution Orbitrap (Thermo Fisher Scientific) coupled with Ultimate 3000 nanoLC (Thermo Fisher Scientific) at Harvard Center for Mass Spectrometry. Peptides were trapped on a trapping cartridge (300 µm x 5 mm PepMap™ Neo C18 Trap Cartridge, Thermo Fisher Scientific) prior to separation on an analytical column (µPAC, C18 pillar surface, 50 cm bed, Thermo Fisher Scientific). Mobile phase system consisted of water with 1% formic acid (A) and acetonitrile with 1% formic acid (B), using a gradient elution for the total of 110 min (1-32% B over 80 min followed by 32-45% B over 20 min), followed by washout with up to 95% B at a flow rate of 300 nl/min. The mass spectrometer operated in positive mode and data dependent acquisition for all analyses. A full scan ranging from 350 to 1400 m/z was performed with a mass resolution of 12×10^4^. The top three most intensive precursor ions from each scan were used for MS2 fragmentation with normalized collision energy of 30 at a mass resolution of 3.0×10^4^. Data were processed using the Proteome Discoverer software (Thermo Fisher Scientific) version 3.1, using a human protein database downloaded in November 2024 from UNIPROT and augmented with the AL-224L sequence. The following criteria were used for identification: cleavage enzyme trypsin (semi-specific); cysteines carbamidomethylation (as a static modification); and methionine oxidation and conversion of N-terminal glutamine into pyroglutamate (as dynamic modifications).

### Short chain fatty acid (SCFA) quantification in fibril extracts

Fibril extracts were tested for the presence of SCFA (1-6 carbons), which were quantified using an ELISA kit (AFG Scientific, Cat. #EK700010) following manufacturer’s protocols. Homogenized and collagenase-treated cardiac tissue (220 mg) was extracted into ten 1 ml fractions as described above. Fraction 3 was used for cryo-EM analysis and fractions 1 to 5 were used for SCFA measurements. Each fraction was diluted 1:5 in the assay buffer and was assayed using 10 µl/well. The assay was performed in technical and biological duplicates. SCFA concentrations were interpolated from the standard curve (R² = 0.97). The results are reported in Figure S6.

#### Assay Validation

To evaluate the antibody selectivity for SCFA vs. non-esterified fatty acids (NEFA) and potential interference from extracellular matrix components in tissue extracts, the following validation experiments were performed. (i) Fractions 1-5 were analyzed to assess the overall assay performance and the interference from NEFA; (ii) Fraction 3 (Fr 3) was spiked with 12 or 24 µmol/l SCFA to test the signal recovery and interference; (iii) 20 µmol/l SCFA was mixed with 20 µmol/l palmitic acid (a representative NEFA) to probe antibody selectivity; (iv) combined spike tests were performed using Fr3 alone, Fr3 + 20 µmol/l SCFA, Fr3 + 20 µmol/l NEFA, and Fr3 + 20 µmol/l SCFA + 20 µmol/l NEFA. The results, reported in Table S4, indicated minimal interference from tissue components and no cross-reactivity with NEFAs, with high signal recovery ranging from 85 to 95%. These results confirm the specificity and robustness of the ELISA measurements of SCFA in cardiac tissue extracts.

The endogenous SCFA concentration measured by ELISA in fraction 3 was 38.7 ± 1.2 µmol/l (or ∼0.17 µmol/g tissue). Total protein concentration in fraction 3 measured by a colorimetric bicinchoninic acid assay was ∼3 mg/ml. Assuming that most of the protein in fraction 3 was the fibril-forming LC fragment 1-116, an approximate molar ratio of LC to SCFA was 3:1. Assuming that SCFA had unit occupancy in the single-filament structure (acetate in Figure 6C), LC to SCFA molar ratio in this structure was 1:1.

### Amyloid assay using thioflavin T fluorescence

Amyloid content in tissue fractions 1-5 was assessed using thioflavin T (ThT), a diagnostic dye that shows increased fluorescence upon binding to amyloid. Fractions 1-5 were diluted 1:5 with Tris NaCl buffer, pH 7.92, and 10 µM ThT was added. Fluorescence emission was measured within 5 min of ThT addition using a Tecan Infinite M1000 Pro plate reader (excitation wavelength λ_ex_=450 nm, emission wavelength λ_em_=480 nm). The measurements were performed in triplicate and the results are shown in Figure S6.

### Cryo-EM sample preparation

Freshly extracted fibrils (water fraction 3) were deposited on the grids immediately upon isolation. UltrAuFoil R 1.2/1.3 copper 300 mesh grids (Quantifoil, N1-A14nAu30-50) were glow-discharged for 45 sec at 15 mA using a PELCO easiGlow glow discharge system to apply a negative charge to their surface. A 3 ml sample was deposited on the grid surface and immediately blotted for 3 sec with a blot force of 3 and plunge-frozen in liquid ethane using a Vitrobot Mark IV apparatus (Thermo Fisher) set to 100% humidity and 4°C.

### Cryo-EM data collection

A cryo-EM dataset of AL-224L amyloid was collected at the Boston University Cryo-EM Core Facility using a Glacios II at 200 kV equipped with a Falcon 4i direct electron detector, Selectris energy filter, and fringe-free imaging. A dataset of 10,309 exposures was collected using Thermo Fisher EPU in counting mode and recorded in the Electron Event Representation (EER) format using a magnification of 130,000x, pixel size of 0.90 Å, total dose of 40 e^−^Å^−2^, dose rate of 10.82 e^−^ per pixel per second, defocus range of −0.8 to −2.5 μm, and energy filter slit width of 10 eV. The imaging rate was approximately 550 exposures per hour. A multishot imaging strategy was used to collect four exposures per grid hole, utilizing beam image shift to move between acquisition areas. Grid screening of the amyloid sample prior to data collection revealed fibrils more abundant near the edges of the foil holes. To maximize the number of fibrils visible in each exposure, the corner of each of the four acquisition areas was positioned partially over the gold film surface.

### Cryo-EM data processing

The dataset was processed in cryoSPARC v4.5.1^59^. Exposures were imported with an EER upsampling factor of 2 then cropped by one-half back to their original resolution using patch motion correction. The contrast transfer function (CTF) correction was performed with ‘patch CTF estimation’. Poor-quality micrographs were removed, yielding 7,076 high-quality micrographs. Template-free ‘Filament tracer’ and ‘inspect picks’ were used to pick filament segments with a separation distance of 45 Å from a 1000 micrograph subset of the data. Particles were extracted at a box size of 896 px and Fourier-cropped to 448 px using ‘extract mics’ to yield a partial uncleaned particle stack (71,140 particles). A set of 2D templates was generated with ‘2D classification’ and ‘select 2D’. The 2D templates were used with ‘filament tracer’ and ‘inspect picks’ to pick filament segments with a separation distance of 45 Å from the full 7,706 micrograph dataset. Particles were extracted at a box size of 900 px and Fourier-cropped to 450 px using ‘extract mics’ to yield a full uncleaned particle stack (421,024 particles). Particle cleaning was performed primarily using 2D classification. Three rounds of ‘2D classification’ and ‘select 2D’ were performed to remove junk particles and double protofilaments yielding a partially cleaned particle stack (236,744 particles). An initial EM map of single protofilament AL-224L was generated using ‘ab-initio reconstruction’ by asking for an output of three volumes and selecting the best quality volume. An initial refinement was performed with ‘helix refine’ to obtain a particle stack with estimated 3D alignments. The partially cleaned, partially aligned particle stack was extracted again at a smaller box size of 400 px. These particles from a smaller box size were cleaned one last time using ‘2D classification’ and ‘select 2D’ to yield a cleaned particle stack (214,351 particles).

A 3D map of the cleaned particle stack was generated with ‘reconstruct only’ and was used as an input for ‘helix refine’. ‘Helix refine’ was used to generate an initial refined EM map of the single-filament amyloid. In this first ‘helix refine’ job, symmetry searching/enforcement was disabled, and the initial model was lowpass filtered to a resolution of 8 Å. The output volume, which was used as an input volume in the next step, showed bumps that correspond to stacked β-sheets. These β-sheet features enabled the symmetry search utility in the next step to find the correct helical rise/twist values. ‘Helix refine’ was performed again with symmetry search/enforcement enabled to produce a Symmetry Enforced Structure. The initial model was lowpass filtered to 4.5 Å, and the twist and rise parameters were searched over -5.2° to 3.2° and 0.8 Å to 8.8 Å, respectively. Helical symmetry search and enforcement was set to begin at 4 Å to ensure that symmetry averaging was applied only after the β-sheet separation along the fibril z-axis was visible. The symmetry was calculated to have a helical rise of 4.86 Å, which is typical of amyloid fibrils, and a twist of 1.20°. Individual particle CTF and individual particle motion was corrected with ‘reference-based motion correction’. The final EM map was generated with a final round of ‘helix refine’ with symmetry search/enforcement enabled (214,100 particles, 2.92 Å, sharpening B-factor of -25.8).

### Cryo-EM model building

An atomic model for AL-224L amyloid was built and refined against the sharpened EM map. Initially, a single polyalanine chain was built into a central section across the fibril z-axis and manually adjusted using Coot^60^. The individual residues were mutated to correspond to the amino acid sequence of AL-224L determined by mass spectrometry. The amino acid assignment within the EM map was determined by carefully examining the path of the backbone, the position of the well-defined disulfide bond, C22-C91, and the strong EM density for side chains that often corresponds to aromatic residues. Residues 42-58 and 105-216 were not modeled because of weak EM density, suggesting conformational heterogeneity in these segments.

The single molecular layer was duplicated to create a five-layer model. The replicated molecules were rigid fit within the EM map above and below the original strand using the fit-in-map function in ChimeraX^61^. The model was further refined in PHENIX^62^ using phenix.real_space_refine^63^. The β-sheet secondary structure, disulfide bond, and non-crystallographic symmetry were manually defined in PHENIX to help guide the refinement and impose identical conformations across all five molecular layers. The model statistics and map-model fit were iteratively improved with multiple rounds of real-space refinements in PHENIX and manual adjustments in Coot. To prevent the model from getting trapped in the wrong conformation at an early stage, initial real-space refinement runs were performed at lower map weight values, while final real space refinement runs were performed at higher map weight values. The model was validated using the cryo-EM Comprehensive Validation module in PHENIX running MolProbity^64^.

To determine the handedness of the cryo-EM map and model of AL-224L amyloid fibrils, we generated a highly sharpened cryo-EM map (sharpening B-factor of -200) and checked if the orientation of the backbone carbonyl atoms matched the corrugation of the backbone cryo-EM density. This approach was described for amyloid-like FUS fibrils^43^ and is suitable for high-resolution cryo-EM maps. In our structure of AL-224L amyloid, the right-handed map and model had a much better map-model fit than the left-handed map and model. Figures were generated with ChimeraX^61^.

### Validation of additional densities using false discovery rate (FDR) thresholding

To validate the significance of the three orphan densities seen in the cryo-EM map at the internal and external sites, we generated a confidence map using FDR thresholding^37^ with default parameters and without incorporating local resolution and atomic model information. We performed the map generation using the unmasked unsharpened AL-224L cryo-EM map as the input and generated a figure viewed at a threshold of 0.99999999, equivalent to a 0.000001% false discovery rate for visible voxels (Supplemental Fig. 1). The three densities were visible in the map indicating that their probability of being noise was below 0.000001%.

## EXTENDED DATA

**Table S1.** Demographic, clinical and laboratory characteristics of case AL-224 with AL amyloidosis at initial evaluation. Abbreviations: BNP, B-type natriuretic peptide; eGFR, estimated glomerular filtration rate; FLC, free light chain; HLC, heavy light chain; Ig, immunoglobulin; LC, light chain; SIFE, serum immunofixation electrophoresis; UIFE, urine immunofixation electrophoresis.

| Characteristics | AL-224 | Reference range |
| --- | --- | --- |
| ***Demographic features*** |  |  |
| Gender | F |  |
| Age, years | 63 |  |
| Age at death, years | 63 |  |
| Cause of death | sudden death |  |
| ***Echocardiographic features*** |  |  |
| Interventricular septal thickness (mm) | 11 |  |
| Ejection fraction, % | 60 |  |
| ***Laboratory parameters*** |  |  |
| Bone marrow plasma cells, % | n/a |  |
| LC restriction | λ |  |
| SIFE | IgG+ λ FLC |  |
| FLC κ | 9.20 | 3.3–19.4 |
| FLC λ | 175.00 | 5.7–26.3 |
| FLC ratio | 0.05 | 0.26–1.65 |
| Abnormal HLC, isotype | IgG |  |
| IgG HLC κ | 1.6 | 3.84–12.07 |
| IgG HLC λ | 8.49 | 1.91–6.74 |
| IgG HLC ratio | 0.19 | 1.12–3.21 |
| UIFE | λ FLC |  |
| BNP, pg/ml | 126 | 0–176 |
| Troponin I, ng/ml | 0.046 | <0.033 |
| Serum creatinine, mg/dl | 0.91 | 0.7–1.3 |
| eGFR, ml/min/1.73 m<sup>2</sup> | 71 |  |
| 24-h urine protein, mg | 469 |  |
| Alkaline phosphatase, U/l | 127 | 25–100 |
| ***Organ involvement*** |  |  |
| Cardiac | + |  |
| Soft tissue | + |  |
| ***Congo red positive histology*** |  |  |
| Fat pad aspirate | + |  |
| Bone marrow biopsy | + |  |

**Table S2.** Top 30 master proteins identified with at least two unique peptides of heart-tissue extracted fibrils by LC-MS/MS. Abbreviations: PSM, peptide-spectrum match; #AAs, number of amino acids in full-length protein; MW, molecular weight. The proteins are ranked according to the number of PSMs. Fibril-forming protein AL-224L (pink), collagen VI chains a1, a 2 and a 3 (green), and the amyloid signature proteins apoA-IV and apoE (orange) are highlighted. Serum amyloid P-component was detected in low abundance (#39, not shown). The data normalization using normalized spectral abundance factor (NSAF) values demonstrated AL-224L as the top identified protein (data not shown). To calculate the NSAF value for each protein, the spectral counts were divided by the protein length (PSM/#AAs) and the sum of all PSM/#AA values was divided by the number of all identified proteins as previously reported (Neilson KA, Keighley T, Pascovici, D, Cooke B, Haynes PA. (2013) Label-Free Quantitative Shotgun Proteomics Using Normalized Spectral Abundance Factors. Methods Mol Biol. 2013:1002: 205-222).

| Accession | Description | Coverage [%] | # PSMs | # Unique Peptides | # AAs | MW [kDa] |
| --- | --- | --- | --- | --- | --- | --- |
| P12883 | Myosin-7 OS=Homo sapiens<br>OX=9606 GN=MYH7 PE=1 SV=5 | 76 | 841 | 133 | 1935 | 223 |
| P13533 | Myosin-6 OS=Homo sapiens<br>OX=9606 GN=MYH6 PE=1 SV=5 | 50 | 521 | 3 | 1939 | 223.6 |
| AL-224L | Amyloidogenic LC sequence AL-224L OS=Homo sapiens | 74 | 373 | 40 | 218 | 23.3 |
| Q8WZ42 | Titin OS=Homo sapiens OX=9606<br>GN=TTN PE=1 SV=4 | 7 | 257 | 224 | 34350 | 3813.7 |
| P62736 | Actin, aortic smooth muscle<br>OS=Homo sapiens OX=9606<br>GN=ACTA2 PE=1 SV=1 | 70 | 187 | 39 | 377 | 42 |
| P12882 | Myosin-1 OS=Homo sapiens<br>OX=9606 GN=MYH1 PE=1 SV=3 | 25 | 157 | 4 | 1939 | 223 |
| P35609 | Alpha-actinin-2 OS=Homo sapiens<br>OX=9606 GN=ACTN2 PE=1 SV=1 | 44 | 112 | 42 | 894 | 103.8 |
| P60709 | Actin, cytoplasmic 1 OS=Homo sapiens<br>OX=9606 GN=ACTB PE=1 SV=1 | 47 | 110 | 5 | 375 | 41.7 |
| P06576 | ATP synthase subunit beta,<br>mitochondrial OS=Homo sapiens<br>OX=9606 GN=ATP5F1B PE=1 SV=3 | 70 | 99 | 58 | 529 | 56.5 |
| Q14896 | Myosin-binding protein C, cardiac-type<br>OS=Homo sapiens OX=9606<br>GN=MYBPC3 PE=1 SV=4 | 38 | 93 | 64 | 1274 | 140.7 |
| P12111 | Collagen alpha-3(VI) chain<br>OS=Homo sapiens OX=9606<br>GN=COL6A3 PE=1 SV=5 | 16 | 76 | 53 | 3177 | 343.5 |
| P25705 | ATP synthase subunit alpha,<br>mitochondrial OS=Homo sapiens<br>OX=9606 GN=ATP5F1A PE=1 SV=1 | 54 | 63 | 41 | 553 | 59.7 |
| P17540 | Creatine kinase S-type,<br>mitochondrial OS=Homo sapiens<br>OX=9606 GN=CKMT2 PE=1 SV=2 | 39 | 57 | 30 | 419 | 47.5 |
| P04004 | Vitronectin OS=Homo sapiens<br>OX=9606 GN=VTN PE=1 SV=1 | 25 | 52 | 17 | 478 | 54.3 |
| Q08043 | Alpha-actinin-3 OS=Homo sapiens<br>OX=9606 GN=ACTN3 PE=1 SV=2 | 14 | 52 | 3 | 901 | 103.2 |
| P06727 | Apolipoprotein A-IV OS=Homo sapiens<br>OX=9606 GN=APOA4 PE=1 SV=4 | 55 | 45 | 37 | 396 | 45.3 |
| P02511 | Alpha-crystallin B chain OS=Homo sapiens<br>OX=9606 GN=CRYAB PE=1 SV=2 | 65 | 36 | 23 | 175 | 20.1 |
| Q9Y2K3 | Myosin-15 OS=Homo sapiens<br>OX=9606 GN=MYH15 PE=1 SV=6 | 3 | 32 | 2 | 1926 | 222 |
| P12109 | Collagen alpha-1(VI) chain<br>OS=Homo sapiens OX=9606<br>GN=COL6A1 PE=1 SV=3 | 19 | 29 | 20 | 1028 | 108.5 |
| P48735 | Isocitrate dehydrogenase [NADP], mitochondrial OS=Homo sapiens OX=9606 GN=IDH2 PE=1 SV=2 | 33 | 28 | 15 | 452 | 50.9 |
| P07437 | Tubulin beta chain OS=Homo sapiens OX=9606 GN=TUBB PE=1 SV=2 | 34 | 28 | 3 | 444 | 49.6 |
| P68371 | Tubulin beta-4B chain OS=Homo sapiens OX=9606 GN=TUBB4B PE=1 SV=1 | 38 | 28 | 4 | 445 | 49.8 |
| P17661 | Desmin OS=Homo sapiens OX=9606 GN=DES PE=1 SV=3 | 33 | 27 | 15 | 470 | 53.5 |
| P12814 | Alpha-actinin-1 OS=Homo sapiens OX=9606 GN=ACTN1 PE=1 SV=2 | 14 | 27 | 6 | 892 | 103 |
| P55809 | Succinyl-CoA:3-ketoacid coenzyme A transferase 1, mitochondrial OS=Homo sapiens OX=9606 GN=OXCT1 PE=1 SV=1 | 28 | 27 | 19 | 520 | 56.1 |
| P54296 | Myomesin-2 OS=Homo sapiens OX=9606 GN=MYOM2 PE=1 SV=3 | 15 | 23 | 19 | 1465 | 164.8 |
| P21796 | Non-selective voltage-gated ion channel VDAC1 OS=Homo sapiens OX=9606 GN=VDAC1 PE=1 SV=2 | 57 | 22 | 16 | 283 | 30.8 |
| P12110 | Collagen alpha-2(VI) chain OS=Homo sapiens OX=9606 GN=COL6A2 PE=1 SV=4 | 16 | 21 | 18 | 1019 | 108.5 |
| P45880 | Voltage-dependent anion-selective channel protein 2 OS=Homo sapiens OX=9606 GN=VDAC2 PE=1 SV=2 | 34 | 21 | 10 | 294 | 31.5 |
| P02649 | Apolipoprotein E OS=Homo sapiens OX=9606 GN=APOE PE=1 SV=1 | 34 | 21 | 15 | 317 | 36.1 |

**Table S3.** Solvation energy calculated for the available amyloid structures of AL LCs. ΔG values represent free energy change upon interface formation between two fibril layers, which was estimated using Protein Interfaces, Surfaces and Assemblies (PDBePISA) server, https://www.ebi.ac.uk/msd-srv/prot_int/cgi-bin/piserver. Bottom two rows list hypothetical values for the λ6-LC germline sequence placed in the amyloid structure of cardiac AL55 LC (PDB: 6HUD) or cardiac AL-224L (PDB: 9OKA). Mutations to restore the germline sequence were introduced using the Swiss-PdbViewer software, https://spdbv.unil.ch/ and the structure was energy-minimized to relieve steric clashes.

| LC sub-family | PDB ID | $\Delta G$,<br>kcal/mol |
| --- | --- | --- |
| $\lambda 3$ | 9FAA | -33.7 |
| $\lambda 3$ | 9FAB | -33.4 |
| $\lambda 3$ | 9FAC | -34.2 |
| $\lambda 3$ | 8R47 | -26.4 |
| $\lambda 3$ | 9EME | -25.5 |
| $\lambda 1$ | 7NSL | -26.6 |
| $\lambda 1$ | 6IC3 | -24.2 |
| $\lambda 3$ | 6Z1O | -21.5 |
| $\lambda 3$ | 6Z1I | -20.4 |
| $\lambda 6$ AL55 renal | 8CPE | -23.3 |
| $\lambda 6$ AL55 cardiac | 6HUD | -20.2 |
| **$\lambda 6$ AL-224L cardiac** | **9OKA** | **-18.0** |
| $\lambda 6$ germline | 6HUD | -18.6 |
| $\lambda 6$ germline | 9OKA | -22.1 |

**Table S4.**
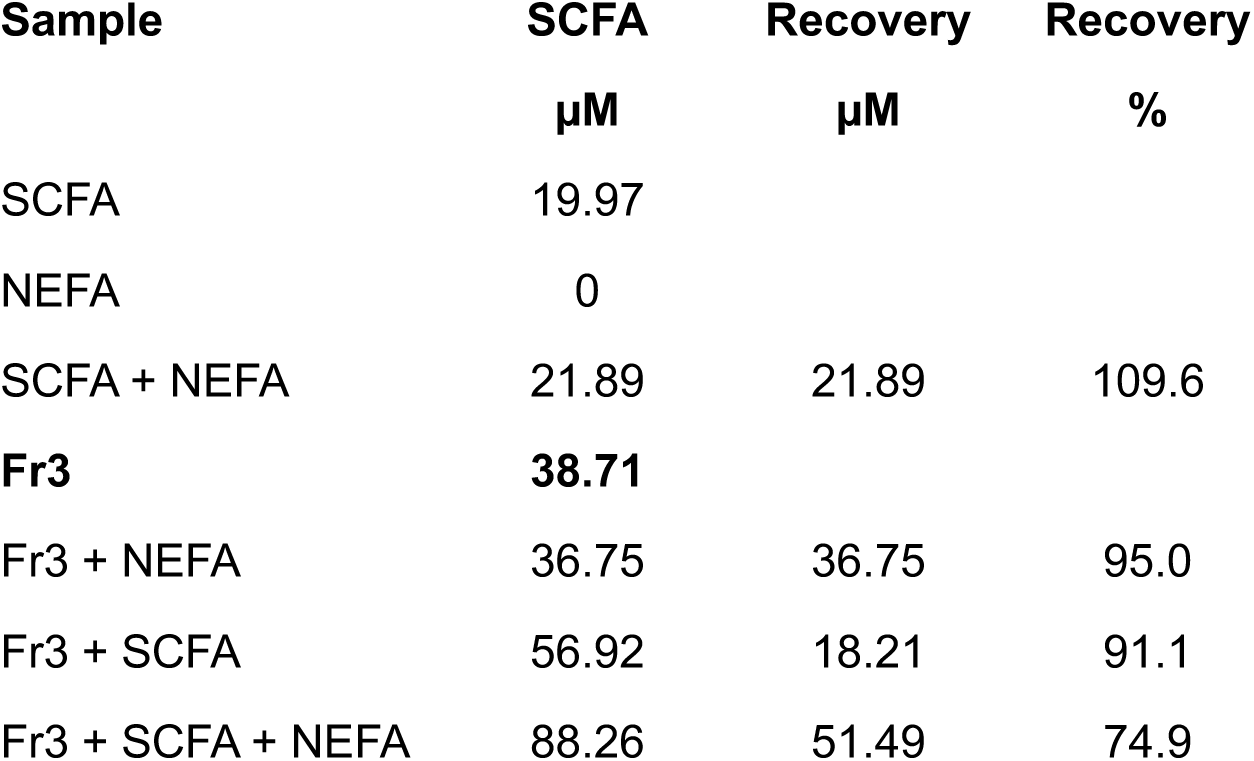
Short chain fatty acid content in the amyloid-containing tissue extracts (fraction 3) measured by ELISA and assay validation. Abbreviations: SCFA - short chain fatty acid; NEFA – non-esterified fatty acid; Fr3 – tissue-extracted fraction 3 used for cryo-EM analysis of amyloid. The experimental details are reported in Methods.

**Figure S1.**
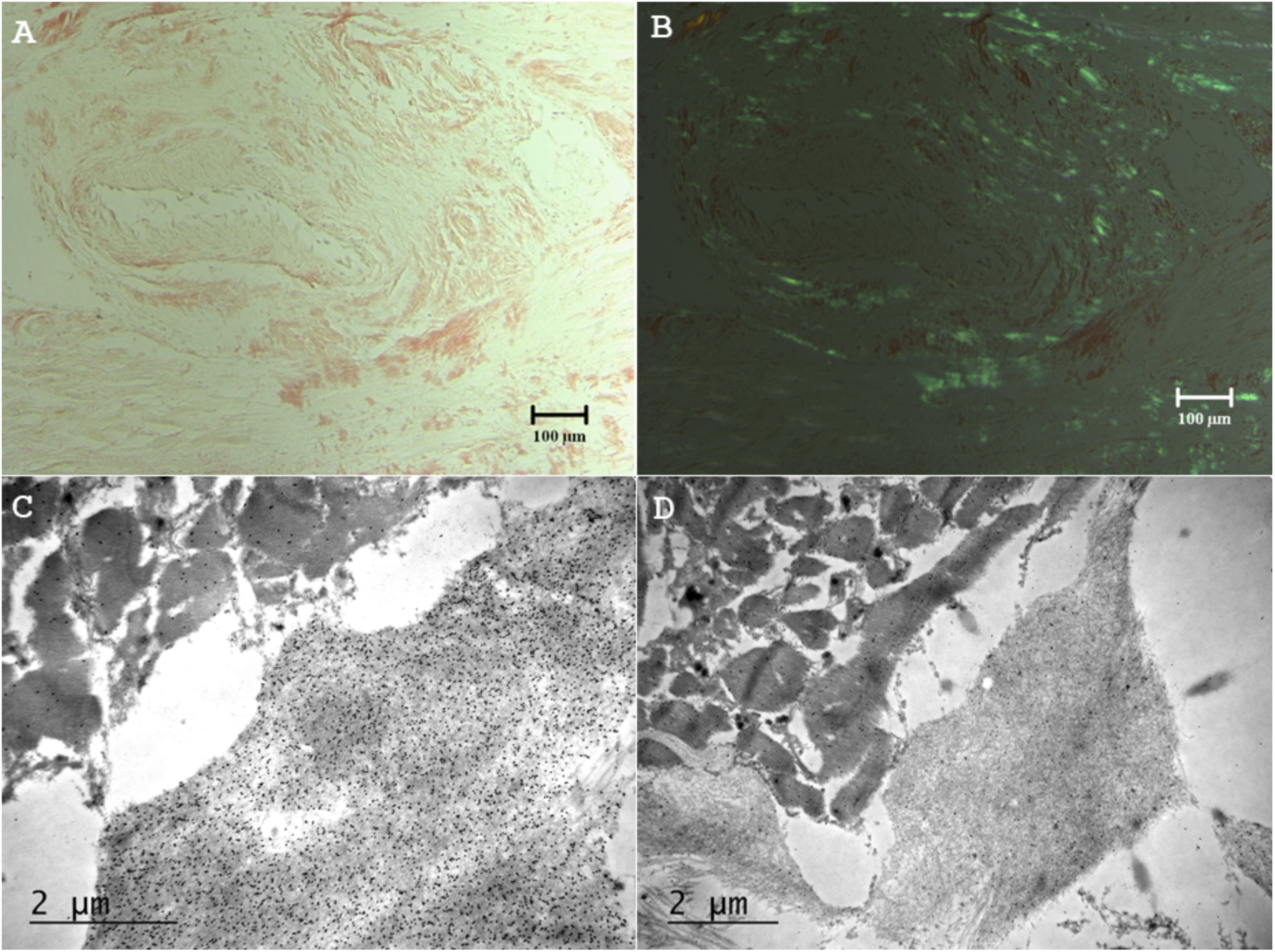
Histological analyses of the autopsied cardiac tissue in case AL-224. **A**, Light microscopy images of Congo red-stained cardiac tissue indicate amyloid deposits. Original magnification ×100. **B**, Areas in panel A viewed by polarized microscopy show amyloid deposits with characteristic green birefringence. Original magnification ×100. **C, D**, Electron micrographs of autopsied cardiac tissue show haystack-like organization of amyloid fibrils in the extracellular space. Immunogold labeling demonstrates numerous electron dense deposits with antibody directed against immunoglobulin λ-LC (original magnification ×20,000, **(C)** and no immunoreactivity with antibody directed against immunoglobulin κ-LC (original magnification ×15,000, **(D)**.

**Figure S2.**
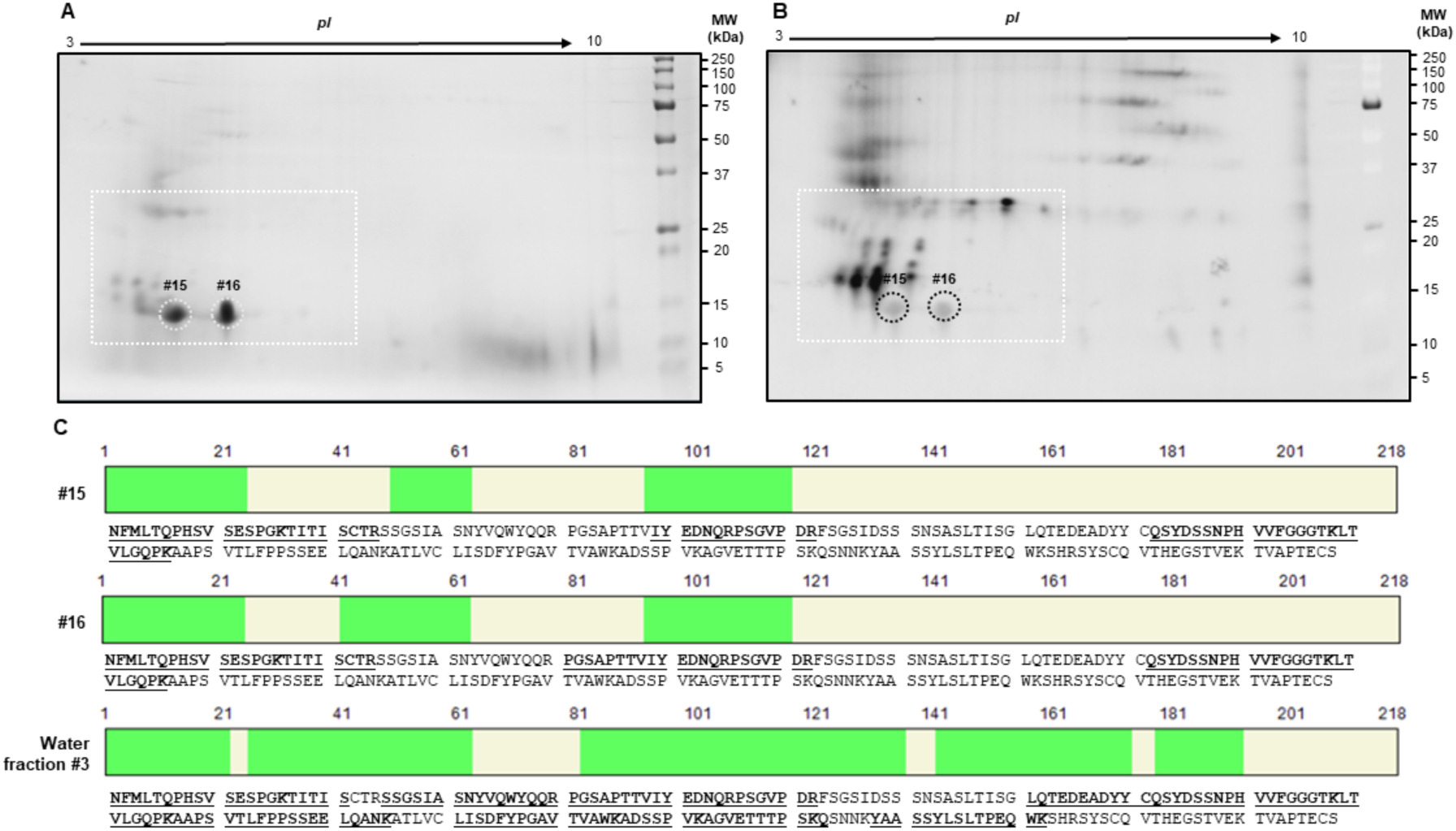
Analysis of AL-224 proteins in water fraction 3 of cardiac tissue fibril extract using 2D SDS-PAGE, 2D western blotting and LC-MS/MS. **A**, Coomassie-stained gel and **B**, the corresponding 2D western blot probed with a primary anti-human λ light chains antibody. Two most prominent spots are circled and marked #15 and #16. These spots were excised and analyzed by LC-MS/MS, along with the entire water fraction 3. **C**, Amino acid sequence of AL-224L shows the position (in green) and sequence (underlined bold) of the peptides identified in spots #15, #16 and in the entire water fraction 3 (indicated).

**Figure S3.**
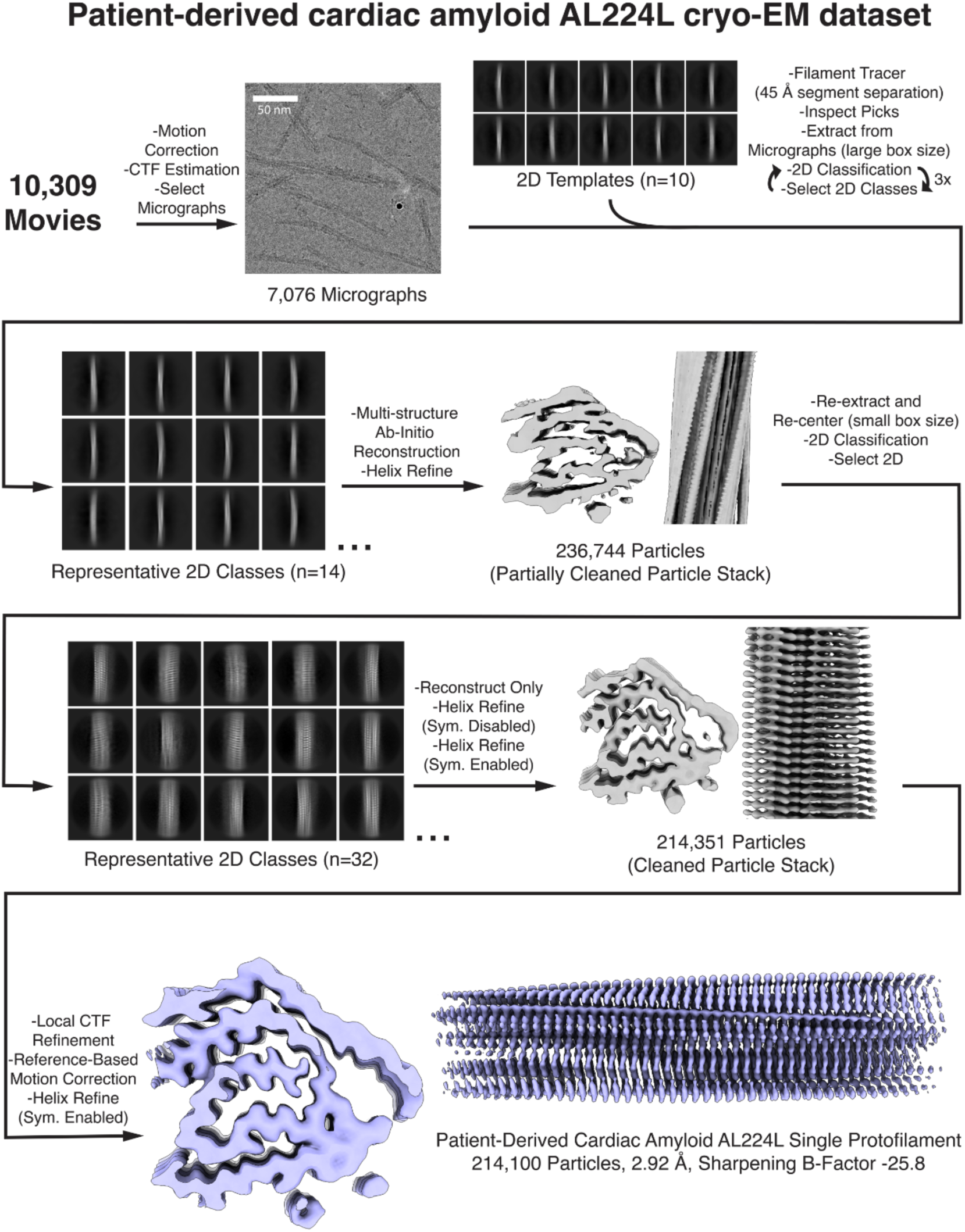
Cryo-EM data processing workflow. General processing pipeline to generate an EM map of an AL-224L single-filament amyloid polymorph.

**Figure S4.**
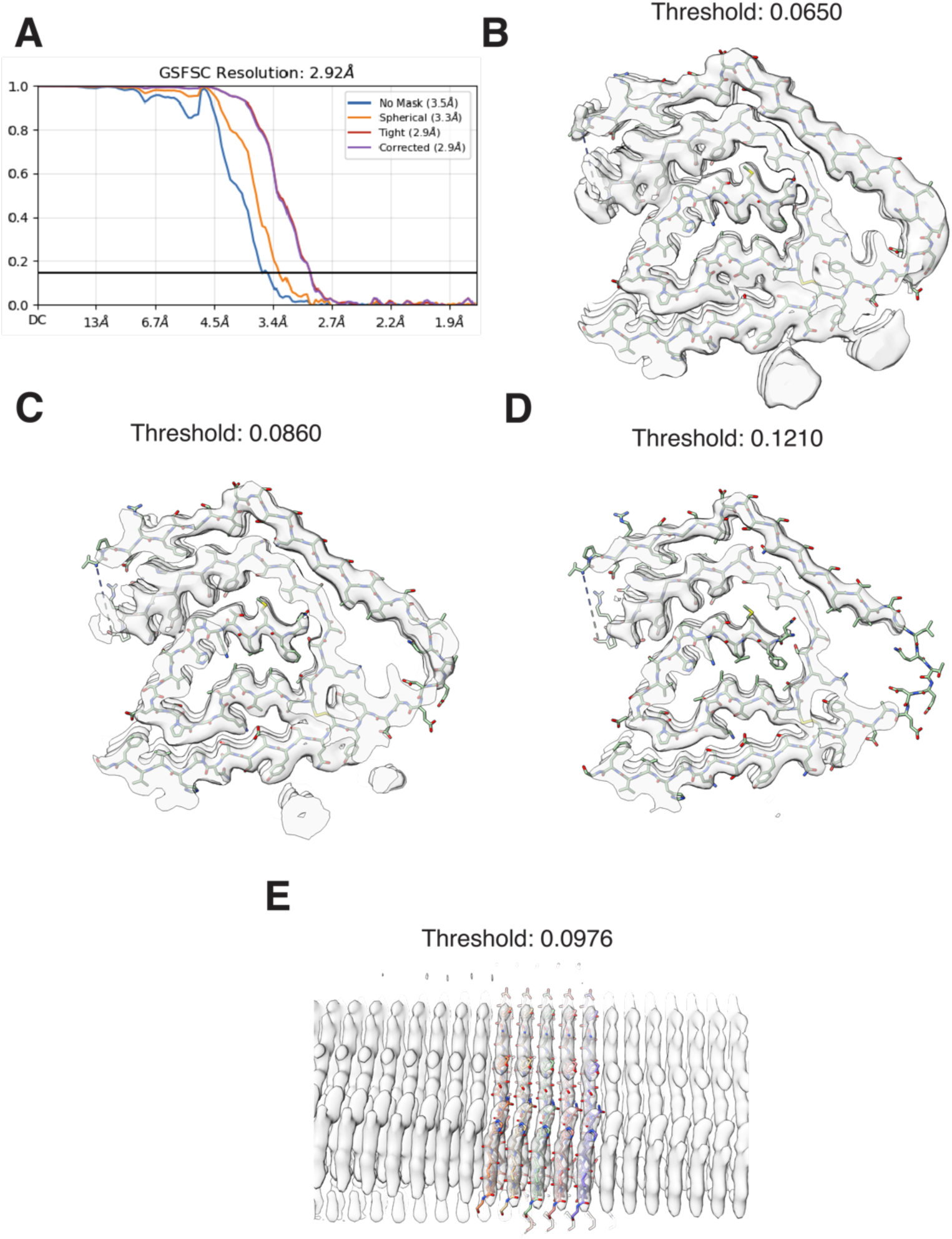
Global resolution and map-model fit evaluation of amyloid structure. **A**, Fourier shell correlation (FSC) plot at a gold standard 0.143 cutoff of cardiac amyloid AL-224L. **B-E**, AL-224L model superimposed over the EM map to show the map-model fit at EM map at thresholds as indicated.

**Figure S5.**
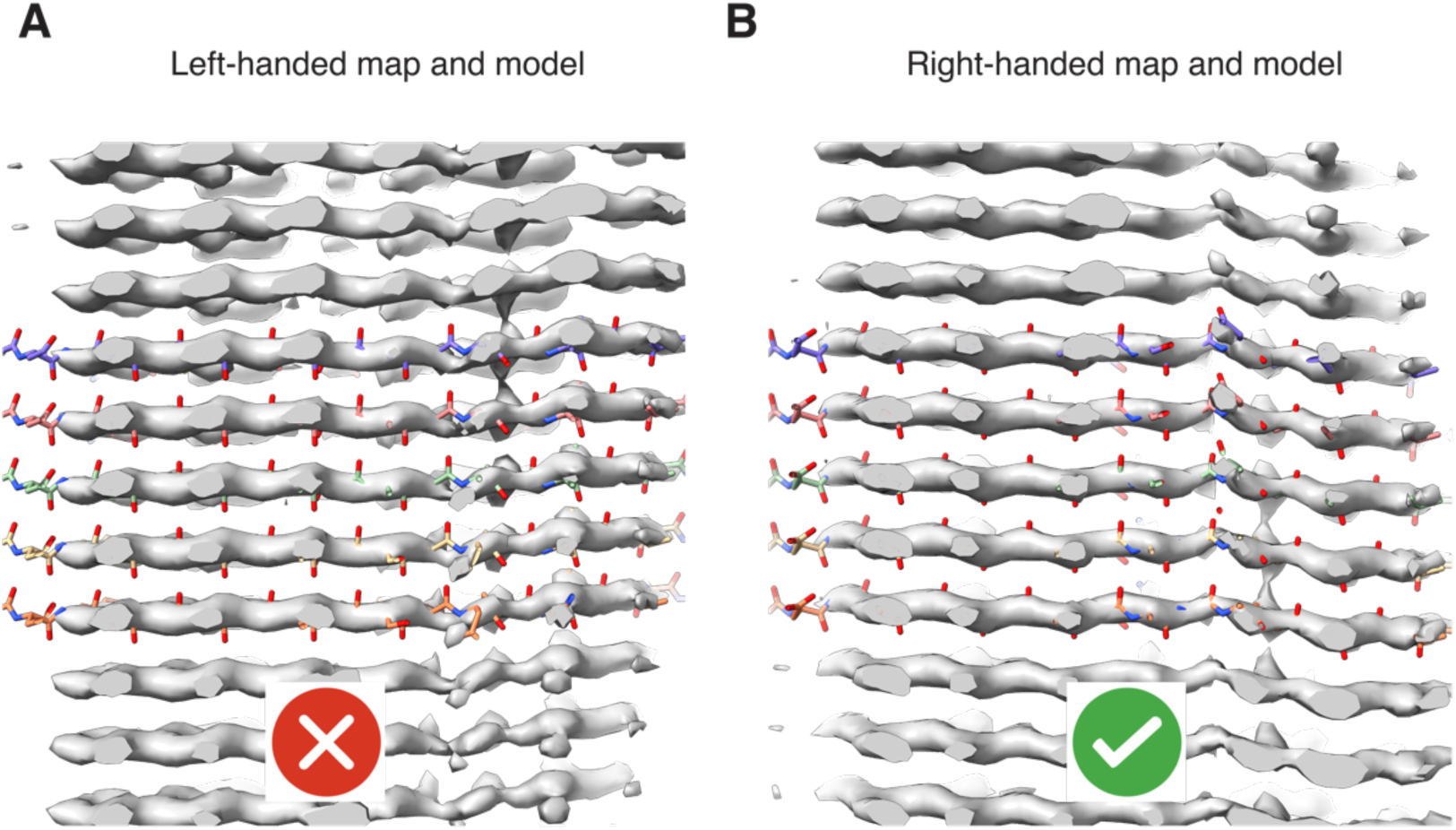
Backbone corrugations reveal the AL-224L amyloid to be right-handed. **A**, Slice-view showing a superimposition of the left-handed amyloid model over the highly sharpened left-handed amyloid map. Green check icon indicates that the right-handed map and model have correct handedness. **B,** Slice-view showing a superimposition of the right-handed amyloid model over the highly sharpened right-handed amyloid map. Green check icon indicates that the right-handed map and model have correct handedness.

**Figure S6.**
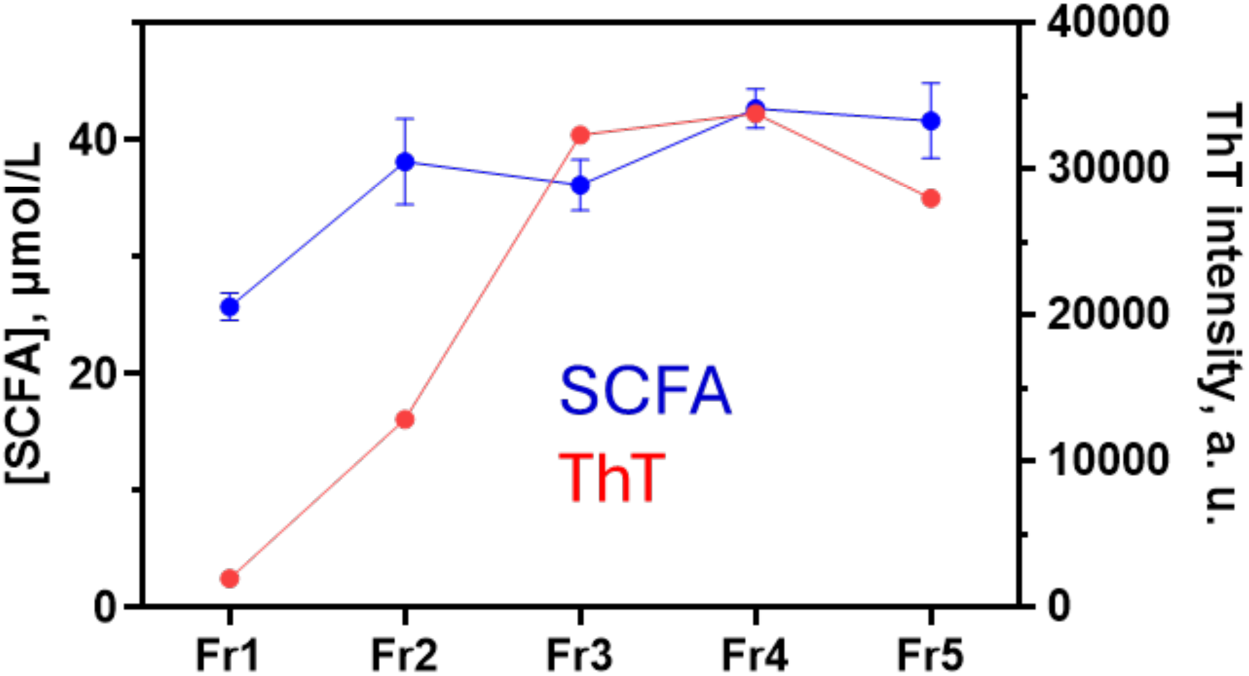
Amyloid and short chain fatty acid (SCFA) content in fractions 1-5 extracted from autopsied cardiac tissue AL-224. SCFA were measured by ELISA in technical and biological duplicates; mean values ± SEM are shown. Amyloid was assessed by measuring fluorescence emission of a diagnostic dye thioflavin T (ThT) that shows increased fluorescence upon binding to amyloid. Methods provide experimental details.

## SUPPLEMENTAL DATA

**Supplemental Figure 1.**
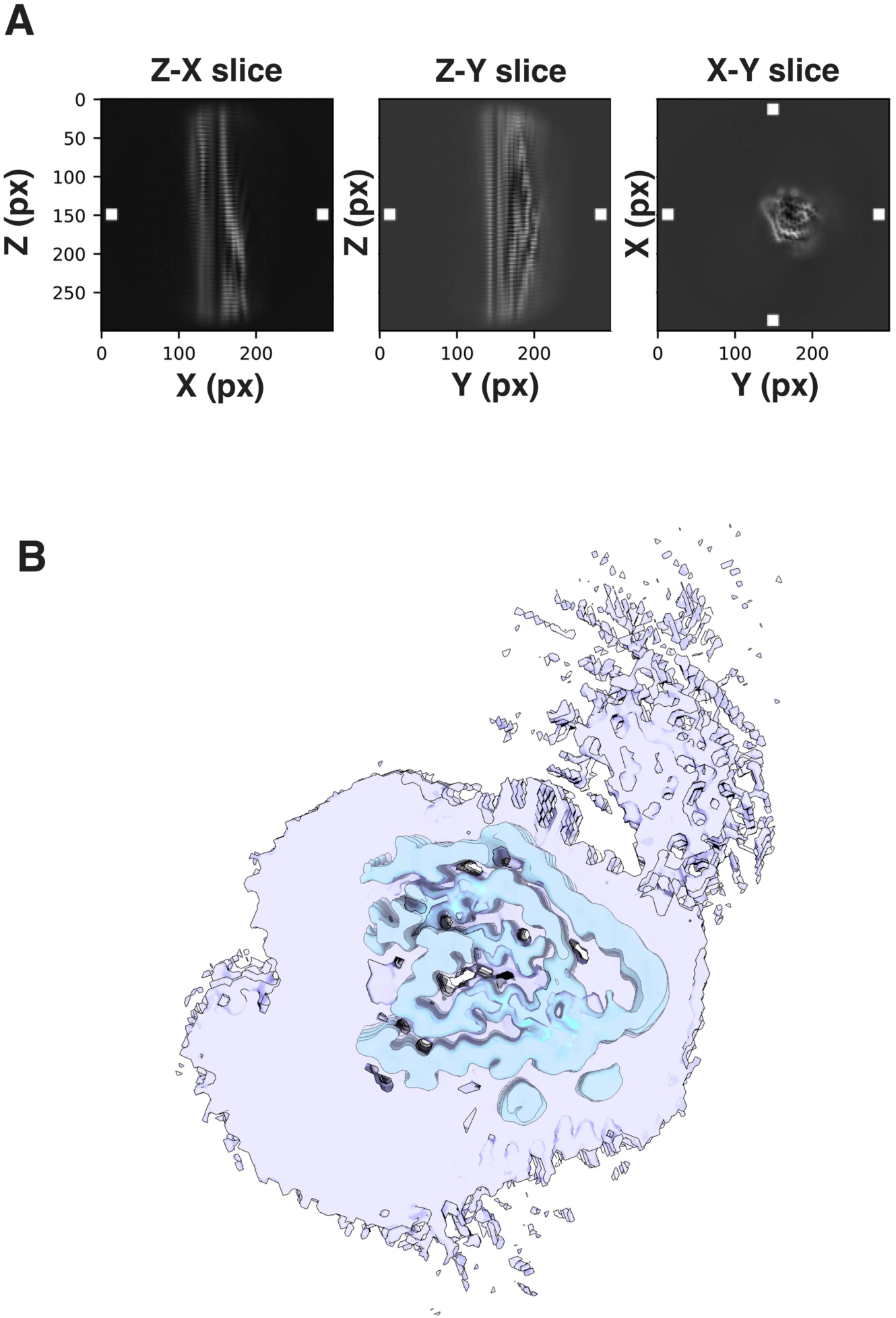
False Discovery Rate (FDR) Threshold Confidence Map. **A**, Slice-views of the unsharpened AL-224L amyloid input EM map to FDR thresholding. White boxes indicate the 3D area that was sampled to produce a noise estimate to generate a confidence map. **B**, Traditional unsharpened AL-224L amyloid EM map (blue) within an unsharpened AL-224L amyloid EM confidence map (purple) generated with FDR thresholding and viewed at a threshold of 0.99999999, equivalent to a 0.000001% false discovery rate for visible voxels.

